# High resolution interaction surface mapping by PRISMA reveals novel ARID1A interactions

**DOI:** 10.64898/2026.03.15.711656

**Authors:** Mercedes Pardo, Chiara Marcozzi, Karen A. Lane, Fernando Sialana, Liudmila Shcherbakova, Zuza Kozik, Mandy Wan, Fuzhou Ye, Claudio Alfieri, Jessica A. Downs, Jyoti S. Choudhary

**Author notes:** Corresponding authors: M.P.,; J.S.C. These authors contributed equally (Chiara Marcozzi, Karen A. Lane). CABIMER, Universidad de Sevilla-CSIC, Spain. The ERC (Epi)Genetic Vulnerabilities in Solid Tumors and Sarcoma Laboratory, Inserm Unit UMR981, Gustave Roussy, and Faculté de Médecine, Université Paris-Saclay, Le Kremlin Bicêtre, France.

## Abstract

The SWI/SNF chromatin remodelling complex controls proliferation and cell fate determination by regulating chromatin accessibility at promoters and enhancers, thereby modulating programs of gene expression, and has roles in DNA damage response, replication, splicing, and translation, and cell plasticity. The cBAF-exclusive subunit ARID1A acts as scaffold for the assembly of cBAF SWI/SNF complexes through its C-terminal globular domain and is the most frequently mutated SWI/SNF subunit in cancer. More than half of the ARID1A protein sequence contains regions of intrinsic disorder which are important for protein interactions, often mediated by short linear motifs. However, these interactions are notoriously difficult to study. Whilst hundreds of ARID1A interactions have been reported in the literature, their molecular basis remains obscure, and only a few have been explored functionally or mapped at an interface level. Here, we use a PRotein Interaction Screen on a peptide MAtrix (PRISMA) combined with quantitative mass spectrometry to identify novel ARID1A interactions and map amino acid residues and motifs that mediate interactions at high sequence resolution. The ARID1A PRISMA assay recapitulates binding of BAF subunits to ARID1A and detects the previously described binding of YAP1 transcriptional coactivator to a PPXY motif. Our PRISMA data reveals binding sites for transcriptional repressor SIN3A and identifies TOX4, CDK2 and CCNA2 as novel interactors. Mutation of a cell cycle-dependent CDK2 phosphorylation site in ARID1A leads to altered gene expression of microtubule factors and defects in cell proliferation. Our work underscores the utility of PRISMA to uncover weak or low abundance interactions that are not detectable by traditional affinity purification strategies. Together, our results characterise novel interactors and a new mode of regulation of ARID1A, and provide a useful resource to further explore mechanistic aspects of ARID1A function.

## Introduction

ARID1A is a defining subunit of the multi-protein cBAF complex, one of several types of SWI/SNF chromatin remodelling complexes that use energy derived from ATP hydrolysis to alter the position of nucleosomes and modify DNA accessibility for RNA polymerase II and other transcriptional regulators, thereby regulating many DNA-based processes, such as transcription, splicing, DNA replication and DNA damage repair ^1–7^. ARID1A directly binds to many of the subunits of the BAF complex and in fact acts as a scaffold for its assembly ^8,9^. Loss of ARID1A leads to complex dissociation, reduced levels of other BAF subunits and BAF complex deficiency ^10–12^. ARID1A has also shown to interact with ribosomes and be important for translation ^13,14^ and reported to have a role in cell plasticity, through a mechano-regulated interaction with transcription factors in the Hippo signalling pathway ^15,16^. Subsequently, ARID1A and BAF function impact many key aspects of cell and organismal physiology ^17,18^. Mutations in ARID1A lead to neurodegenerative diseases and account for ∼6% of all cancer mutations ^19–22^. Better understanding of regulatory mechanisms of ARID1A function could help develop novel therapeutic strategies.

ARID1A is a large protein containing two conserved domains of globular structure, an ARID domain located centrally that mediates binding to DNA and a C-terminal domain that mediates interactions with other BAF subunits and is required for BAF complex formation. In contrast, the N-terminal half of the protein is mostly made up of intrinsically disordered regions (IDRs) ^23^. Not much is known about the function of this region of ARID1A. Despite their lack of structure, IDRs play crucial roles in the control of signalling pathways and can also work as platforms for the assembly of large proteinaceous structures. IDRs frequently contain SLiMs or short linear motifs that mediate interactions with globular domains ^24^. They also often carry post translational modifications (PTMs) that function as regulatory switches ^25^. Many disordered regions are capable of phase separation, and this is indeed a general property of most SWI/SNF subunits ^26^. The N-terminus of ARID1A can partition into phase-separated condensates, possibly through interaction with FUS, a nucleic acid binding protein that forms oncogenic fusion proteins ^26,27^. Recently, the IDR of ARID1A was also shown to be required for genomic targeting of the BAF complex to enhancers and interaction with transcription factors and other transcriptional machinery ^28^.

The majority of ARID1A interacting proteins that have been mapped to date bind to the C-terminal half of ARID1A (Figure 1A). This includes, in addition to other BAF subunits, proteins involved in DNA damage response and repair such as ATR and MSH2, cell cycle regulators such as p53 and RB, and other transcriptional regulators such as Polycomb protein EZH2 and the glucocorticoid receptor, transcription factor HIC2, topoisomerase enzyme TOP2A, and E3 ubiquitin ligase bTrCP. Characterised binding partners on the disordered N-terminal half of ARID1A include FUS, and YAP1/TAZ, transcription factors that regulate cell plasticity ^15^.

**Figure 1.**
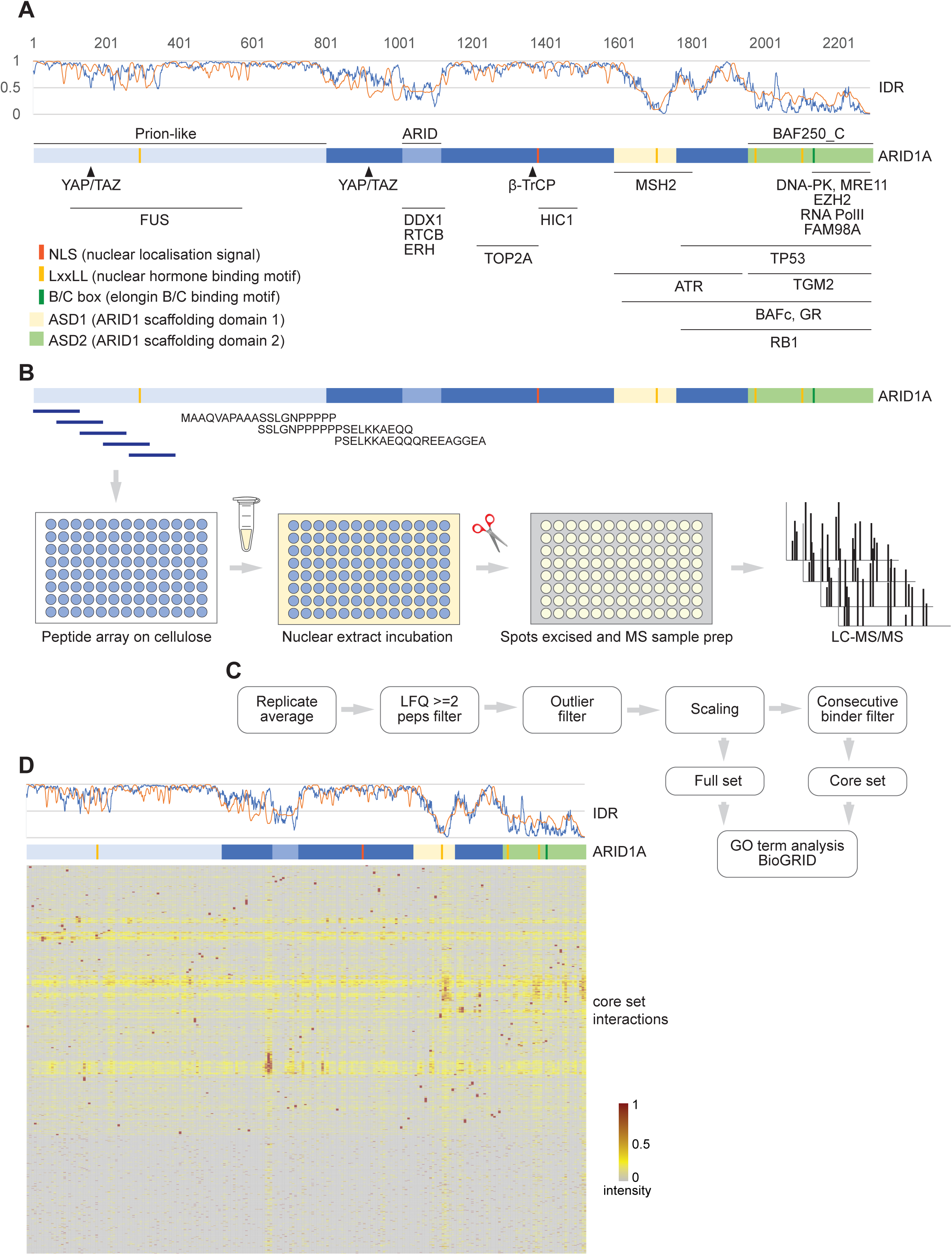
ARID1A PRISMA interactome. **A,** Curated ARID1A physical interaction map. Coloured boxes in ARID1A diagram represent significant domains, regions of motifs. The top graph represents intrinsic disorder regions (IDR). Intrinsic disorder scores were calculated with IUPred2A (https://iupred2a.elte.hu/). The blue line represents the IUPred2 prediction (disordered protein regions) and the orange line represents the ANCHOR2 prediction (disordered binding regions). Scores above 0.5 represent intrinsic disorder. **B,** PRISMA experimental workflow. Nuclear lysates from RPE1 cells were incubated with an ARID1A tiled peptide array. After membrane washes the individual peptide spots containing binding proteins were processed and analysed by mass spectrometry. **C,** PRISMA data analysis workflow**. D,** Clustered heat-map of PRISMA binding profiles aligned with the ARID1A protein diagram and IUPred2A predictions, representing data from a single biological replicate with technical replicate MS measurements for 50% of peptide spots. Only proteins that were identified by at least two peptides were used, and the two replicate datasets were integrated by calculating the average LFQ intensity (hereafter intensity) for each protein in each spot where two measurements were available or using the single available value. The signal intensity for each protein across the 228 bait peptides was then normalised between 0 and 1. See also Tables S1 and S2 and Figures S1 and S2.

In addition to these, many other ARID1A-interacting proteins have been reported in the literature. However, the molecular basis for these interactions is unknown. Recent reports suggest that many functional ARID1A interactions occur through the disordered regions. IDR-mediated interactions are inherently challenging to study due to their reversible and weak nature, and traditional approaches are not best suited to investigate them ^29,30^. Here, we take advantage of a peptide interactomics screen on a tiled peptide array to systematically identify amino acid residues, domains and motifs that mediate ARID1A interactions with other proteins at high sequence resolution. Our PRISMA data recapitulates interaction surfaces of ARID1A with other BAF complex subunits, and SLiM-mediated interactions. We identify and validate several novel ARID1A-interacting proteins with roles in transcriptional regulation and cell cycle control, and map their binding sites. Finally, we functionally characterise and uncover a role for ARID1A phosphorylation in the transcriptional regulation of microtubule organisation and cell division, providing the first insights into ARID1A regulation through phosphorylation.

## Results

### A sequence-resolution biophysical interaction map of ARID1A

To systematically map the landscape of ARID1A protein-protein interactions at high sequence resolution and delineate interaction surfaces, domains and motifs in ARID1A, we took advantage of the recently developed *Protein Interaction Screen on peptide MAtrices* (PRISMA) technology ^31^. The PRISMA assay consists of high-throughput affinity purifications using hundreds of overlapping peptides spanning the entire sequence of a target protein immobilised on a nitrocellulose membrane, combined with shotgun mass spectrometry analysis, for identification and quantitation of peptide-bound proteins. We designed our PRISMA assay with 20-mer peptides with a 10-aa (amino acid) overlap covering the whole length of human ARID1A (2285 aas), resulting in a high-density tiled array of 228 peptide spots (Figure 1B, Table S1). Binding proteins were screened using nuclear extract from RPE1 cells in the presence of nuclease to eliminate DNA-mediated interactions. A total of 2225 proteins identified by at least 2 peptides were quantified (Figure 1C, D).

Around a quarter of the proteins bound ubiquitously across the peptide array at low intensities, while the rest displayed much higher signals in discrete regions of the array. The frequency of binding was calculated for each spot, revealing that the number of interacting proteins per spot ranged from 1 to 235 (Figure 1D). Approximately 700 proteins (30%) were quantified in more than half of the peptide array (100 peptide spots), whilst around 100 were found to bind to over 200 peptide spots (Figure 1D). This so called “promiscuous” binding has also previously been observed in a PRISMA assay for C/EBPβ ^31^. To reduce the noise from background binding and highlight the strongest signals across the peptide array, binding signals below the 75^th^ percentile intensity were removed. Furthermore, to discriminate robust interactions, a second filter based on the presence of consecutive signal intensities across a minimum of two contiguous peptides was applied. The consecutive binding is driven by the overlap in the tiled peptide sequences (Figure 1B, C). This strict filter criterion, applied after the 75^th^ percentile outlier filtering, yielded an ARID1A core PRISMA interaction set of 1454 proteins. It is worth pointing out that the consecutive binding filter when applied to the ARID1A PRISMA dataset is much more restrictive that the one applied in the original PRISMA paper, where the matrix peptides were shorter (11 aas) and also had a smaller overlap (4 aas), which means that generally any sequence stretch was covered by three peptides, whilst in the ARID1A PRISMA assay described here any sequence is only covered by two peptides. The distribution of peptide interactions was not attributed to physicochemical parameters of the peptides, as evidenced by comparison of the number of binding proteins with peptide properties such as hydrophobicity, charge, and isoelectric point (Figure S1). These results suggest that the PRISMA peptides on the matrix retained their specific protein-binding properties.

The global binding profiles from the PRISMA dataset were represented as a hierarchically clustered heatmap of normalised intensities (Figure 1D, Figure S1, Table S2). This revealed there were large differences in the binding profiles of the matrix peptides indicating that binding was specific, as background binding would be expected to result in a homogeneous distribution of interacting proteins. Peptides encompassing the conserved <u>A</u>RID <u>s</u>caffolding <u>d</u>omains ASD1 and ASD2, and particularly those containing the LXXLL motifs #2 (aas 1709-1713) and #4 (aas 2085-2089), displayed the highest number of interactions, followed by regions flanking the ARID domain (Figure S1). The ARID DNA-binding domain and the NLS appeared devoid of interactions. The region immediately upstream of the ARID domain that sustains a high number of interactions is an unstructured stretch that is predicted to fold upon binding to a protein partner by the ANCHOR2 algorithm ^32^. The N-terminal region also displayed significant numbers of interactions, seemingly organised in locally enriched hotspots, some of which overlapped with predicted disordered sequences or disordered binding regions.

GO term enrichment analysis of the PRISMA core interaction dataset revealed significant over-representation of proteins annotated with GO terms related to known ARID1A functions (Figure S2A, Table S3), with enriched terms including translation, mRNA splicing, ATP-dependent chromatin remodelling and DNA damage response. Enrichment of protein domains revealed that nucleotide binding domain, RNA recognition domain, Armadillo, DNA/RNA helicase domain and AAA ATPase domain were some of the most frequently represented domains and sites (data not shown). To further assess the biological relevance of the PRISMA-derived data, we compared it to the ARID1A physical interactome retrieved from the BioGRID repository of protein interactions. Interactors identified by yeast two-hybrid or RNA-purification, viral proteins and duplicates were removed prior to comparison, leaving 211 ARID1A physical interactors. Forty-nine of these were detected by PRISMA and 31 of them were included in the core dataset (Figure S2B). These results suggest that the PRISMA-derived interaction data shows a high level of overlap and coverage of the native ARID1A interactome and represents a useful resource to explore sequence-to-function relationships at high sequence resolution.

### PRISMA recapitulates binding of BAF subunits

To assess the ability of the ARID1A PRISMA to detect localised interactions, we investigated the binding of components of the BAF complex. Core module subunits SMARCB1, SMARCC1/2, SMARCD1/3 and SMARCE1, ATPse module subunits SMARCA4 and ACTL6A, and accessory subunit DPF2 displayed correlated interaction footprints on the conserved ASD1 and ASD2 domains, immediately upstream of the ARID domain, and on the central part of the disordered N-terminal domain, with the latter displaying the strongest signals (Figure 2A). Consecutive binding signals of SMARCA4, SMARCB1 and SMARCE1 were observed in the ASD2 domain aligned with the LXXLL motif located at position 2085-2089, which has been shown to be essential for binding of BAF subunits ^33^. We then compared binding sites of BAF subunits with cryogenic electron microscopy (cryoEM) structural model 6LTJ data ^9^. The PRISMA interaction footprint was in good agreement with cryoEM maps, demonstrating multiple contacts of the ASD1 and ASD2 conserved domains of ARID1A with SMARCA4, SMARCB1, SMARCC1/2, SMARCD1, SMARCE1 and DPF2 (Figure 2B). Furthermore, binding signals of SMARCA4, SMARCB1 and SMARCE1 showed remarkable overlap with protein contact sites from the cryoEM structural models (Figure 2B). In this cryoEM study only a stretch of the ARID1A C-terminus (aas 1639-2285) was modelled, precluding the comparison of contact sites at the protein N-terminal half. Therefore, we also compared the PRISMA data to BAF complex crosslinking-mass spectrometry (XL-MS) data ^8^ that encompasses the whole ARID1A sequence. The agreement between both was particularly noteworthy (Figure 2B): in addition to the well-documented binding of several BAF subunits on the C-terminal region of ARID1A, both PRISMA and XL-MS showed SMARCD, SMARCC and ACTL6A strongly binding to ARID1A N-terminal region. The overlap was observed in regions, not in exact positions, as the different approaches provide different resolution and measure different properties, but converged on a consistent structure. Hence, the PRISMA data mirrored binding data from other experimental approaches and provided information on contact sites between associated proteins. Our PRISMA data suggests that while the BAF250_C region is essential for scaffolding the assembly of the BAF complex ^33^, the disordered N-terminus also mediates contacts with different BAF subunits, in agreement with XL-MS ^8^. Most SWI/SNF subunits have disordered prion-like regions that are involved in multivalent protein interactions ^23^. Interactions engaged by these disordered regions may serve a regulatory function.

**Figure 2.**
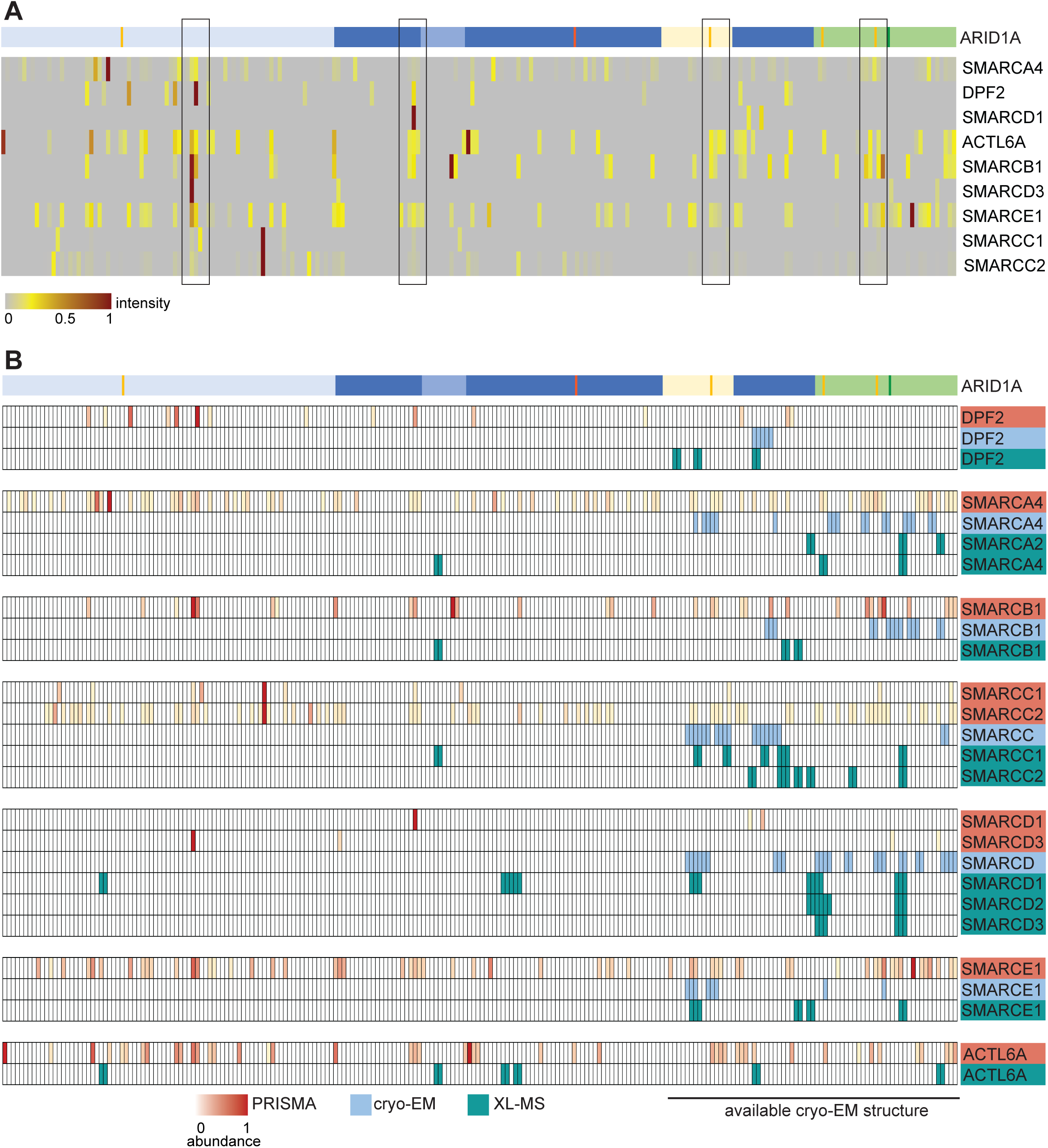
PRISMA captures ARID1A binding interfaces to BAF subunits. **A,** ARID1A PRISMA profiles of BAF subunits**. B,** Detailed comparison of PRISMA data on BAF subunits to amino acid contacts in cryoEM model structure 6LTH ^9^ and XL-MS data of the SWI/SNF complex ^8^. No contacts between ARID1A and ACTL6A were observed in cryoEM structure 6LTH. Contacts by different paralog subunits are represented where available.

Next, the PRISMA data was compared with published sites of interaction in ARID1A with other proteins (Figure 3A). Out of 16 mapped ARID1A direct binders found in the PRISMA data, three displayed PRISMA binding patterns that agreed with their mapped site of interaction with ARID1A. Transcriptional coactivator YAP1 displayed consecutive binding at 3 peptides spanning ARID1A aas 131-170 (Figure 3B). These peptides contain the proline-rich PPXY sequence, a SLiM that mediates binding of ARID1A to the YAP1 WW domain, and mutation of ARID1A Tyr148 to Ala has been shown to eliminate YAP1 binding ^15^. Therefore, our PRISMA data successfully pinpointed the site of YAP1 interaction. Other WW domain proteins such as ITCH and NEDD4L were also detected binding to these PPXY containing peptides in our data, indicating that PRISMA can truly recapitulate SLiM-mediated direct associations. In contrast, transcriptional coactivator TAZ, which belongs to the same family as YAP, displayed consecutive binding at peptides from the ARID1A C-terminus and semi-consecutive binding between aas 2061-2100. TAZ only contains one WW domain in contrast to YAP1 which contains 2, so this might explain the lack of detection of TAZ at the 144-148 PPXY motif. The retinoblastoma protein (RB1) displayed weak consecutive binding signals on the two most C-terminal peptides (position 2261-2285), in agreement with published data ^34^. ARID1A has been shown to interact with Rb and suppress its phosphorylation, with two truncation fragments, ARID1A:1110-1976 and ARID1A:1977-2285, being able to mediate the interaction. The binding footprint of paraspeckle protein FUS showed strongest signals in the disordered N-terminus of ARID1A, and some residual binding in a disordered region upstream of the ASD1 domain. This is also consistent with published data showing that ARID1A interacts and partitions into biocondensates with FUS through its N-terminal prion-like domain ^26,35^. Other mapped ARID1A binders displayed only scattered single signals that could not be confidently attributed to specific binding (TOP2A, MSH2, ATR) or showed promiscuous binding (TGM2, ERH and PRKDC).

**Figure 3.**
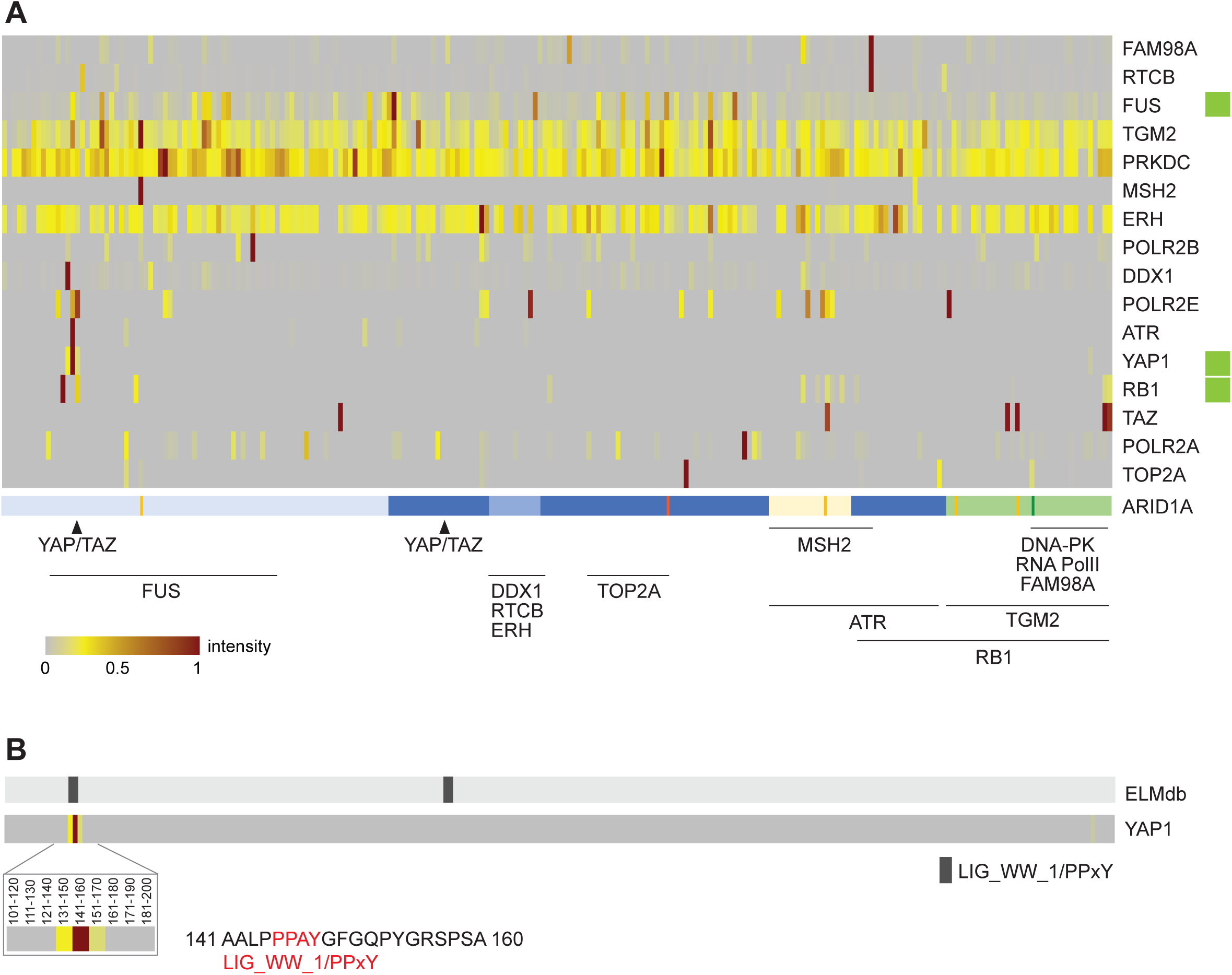
PRISMA refines interaction interface maps of known ARID1A interactions. **A,** PRISMA profiles of ARID1A mapped interacting partners**. B,** PRISMA detects YAP1 binding to a PPXY SLiM in ARID1A. ELMdb represents Eukaryotic Linear Motifs database annotation of PPxY motifs (LIG_WW_1) on ARID1A. Intensity scale as in A.

In summary, our results demonstrate that the ARID1A PRISMA assay can recapitulate FUS, RB1 and YAP1 in vivo binding. These have not previously been identified in affinity purification-mass spectrometry (AP-MS) interactomes, suggesting that the *in vitro* set-up of PRISMA may be able to reveal complementary interactions that might be too labile or of too low abundance for traditional AP-MS approaches.

### PRISMA reveals binding of SIN3A to a predicted SIN3A SLiM

Seeing as PRISMA can recapitulate SLiM-mediated interactions, we next investigated whether our assay could identify binding to other ARID1A SLiMs. We used the Eukaryotic Linear Motif database (ELMdb) ^36^ prediction tool to obtain a list of potential SLiMs in human ARID1A and checked whether peptides containing these motifs displayed significant specific binding of proteins containing the matching globular domain. We observed highest binding signal of E3 ubiquitin ligases on peptides containing degrons, and SH3 domain proteins on peptides containing an SH3-binding SLiM (Table S1). Detailed inspection of the data revealed strong consecutive binding of the transcriptional repressor SIN3A to peptides encompassing a predicted Sin3-interacting domain (SID) of low conservation (Figure 4A), suggesting this motif could mediate binding between SIN3A and ARID1A. The PRISMA data also showed SIN3A binding to the two C-terminal peptides (aas 2261-2285). ARID1A and SIN3A have been shown to physically interact to repress the expression of cell cycle genes in mouse osteoblast precursor cells ^37^, but the direct binding determinants have never been mapped. SIN3A has also been shown to interact with other subunits of BAF complexes ^38–41^. Therefore, our results suggest that ARID1A and SIN3A may interact directly through a Sin3 SLiM and other supporting nearby residues.

**Figure 4.**
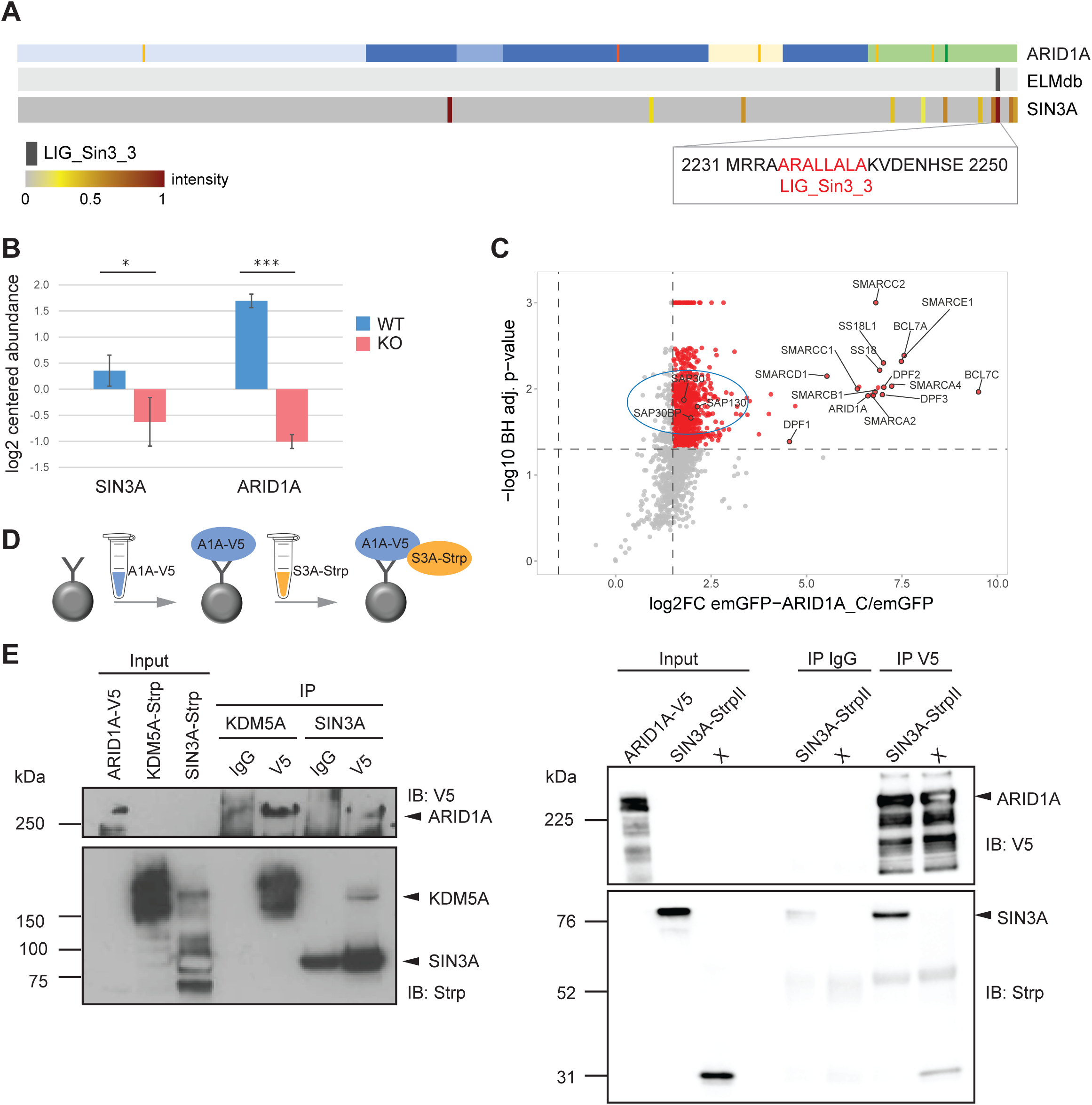
SIN3A binds to ARID1A through a poorly conserved Sin3 SLiM. **A,** Alignment of the SIN3A PRISMA binding profile to ARID1A tiled peptide array and the Sin3 SLiM ELM prediction on ARID1A**. B,** Bar graph representing levels of SIN3A on chromatin in ARID1A KO and parental RPE1 cells (n=3, mean ± SD, * p< 0.05, *** p< 0.00005). **C,** Co-immunoprecipitation of endogenous SIN3 complex (circled) by ARID1A C-terminal fragment GFP pulldown. **D,** Schematic of the *in vitro* binding experiment between recombinant ARID1A-V5 and SIN3A fragment (449-1086) containing the PAH3 domain tagged with StrepII tag. **E,** Co-immunoprecipitation of recombinant ARID1A-V5 and SIN3A_PAH3-StrepII. ARID1A-V5 was immunoprecipitated from insect cell lysates. IgG was used as a negative control. Beads were then incubated with cell lysate from insect cells expressing SIN3A_PAH3-StrepII. Co-immunoprecipitation was assessed with anti-V5 and anti-StrepII antibodies to detect ARID1A and SIN3A_PAH3 respectively. KDM5A-StrepII was used as a positive control. Lanes labelled with X are irrelevant for this work.

ARID1A is required for the recruitment of SIN3A to chromatin in ovarian cancer cells and quiescent mouse pre-osteoblastic MC3T3-E1 cells ^37,42,43^. To assess if ARID1A is necessary for chromatin recruitment of SIN3A in RPE1 cells, we isolated chromatin from ARID1A KO RPE1 cells and their wild type counterparts and performed quantitative profiling of chromatin-binding proteins by mass spectrometry. Inspection of the chromatome revealed that levels of SIN3A on chromatin were significantly lower in ARID1A KO cells compared to WT cells (Figure 4B), confirming that ARID1A is required for the recruitment of SIN3A to chromatin in RPE1 cells.

To assess if the association between ARID1A and SIN3A requires the ARID1A N-terminus, we expressed an ARID1A C-terminal fragment encompassing aas 1018-2285 tagged with Emerald-GFP (Em-ARID1A_C) in ARID1A KO RPE1 cells and performed GFP-trap immunoprecipitation followed by MS analysis. All subunits of the SIN3 complex were enriched in ARID1A_C immunoprecipitates compared to control GFP-only (Figure 4C), confirming that ARID1A C-terminus is sufficient for association with the SIN3 complex.

The SID SLiMs of MAD1 and other proteins specifically interact with the PAH domain of SIN3 ^44^. To confirm if ARID1A and SIN3A interact directly, we individually expressed full length ARID1A fused to the V5 tag ^45^ and a SIN3A fragment encompassing aas 449-1086 which includes the PAH3 domain (SIN3A_PAH3) tagged with StrepII in insect cells and performed pull-down experiments (Figure 4B, C). Full length KDM5A-StrepII was included as a positive control, since its paralog KDM3A has previously been shown to bind to ARID1A ^46^. Bead-bound ARID1A captured KDM5A and SIN3A_PAH3 (Figure 4C). Although SIN3A_PAH3 was also detected in the negative control IgG immunoprecipitated, it was consistently much more abundant in V5 immunoprecipitates. Therefore, our results indicate that ARID1A interacts directly with the PAH domain of SIN3A.

### PRISMA identifies novel ARID1A interactor TOX4

Comparison of PRISMA data to ELM database predictions revealed potential novel candidate interactor TOX4 binding to two ARID1A peptides spanning aas 611-630 and 1691-1710 (Figure 5A). TOX4 is a regulatory subunit of the protein phosphatase 1 (PP1) complex that is almost exclusively associated with chromatin and has been shown to mediate dephosphorylation of the C-terminal domain (CTD) of RNA polymerase II, thus regulating transcriptional elongation ^47^. Despite being distal from each other, both TOX4-binding peptides encompassed phosphorylation consensus sites for several kinases (polo-like kinase 1 (PLK1), Casein kinase I (CK1) and glycogen synthase kinase 3 (GSK3)), suggesting that these were specific binding events. Phosphorylation of ARID1A serine 631 (Ser631) located in the kinase consensus site included in peptide 611-630 has been previously detected (Figure S3, data from phosphositeplus.org ^48^).

**Figure 5.**
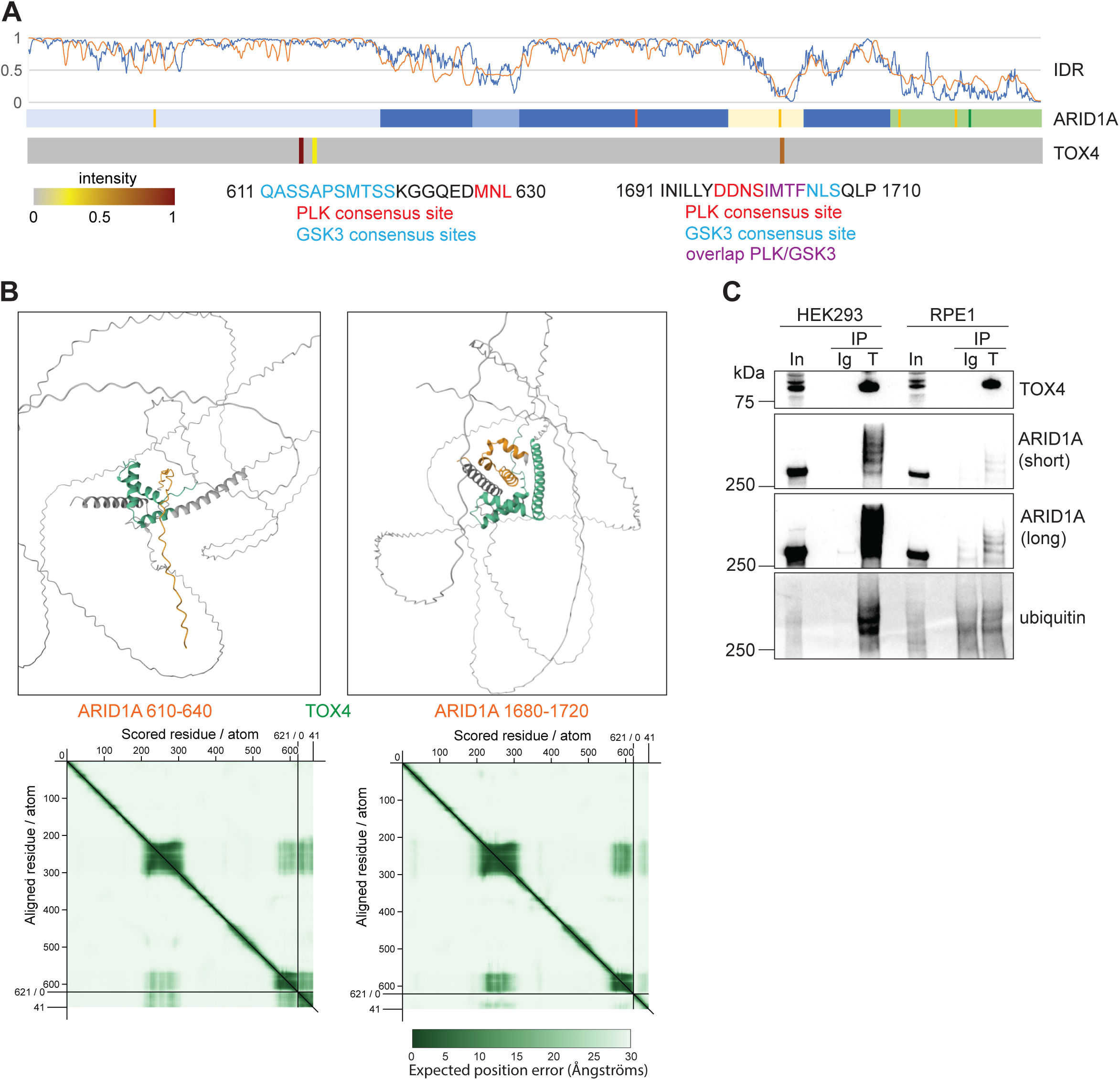
A novel interaction between ARID1A and protein phosphatase 1 regulatory protein TOX4. **A,** TOX4 PRISMA binding profile to ARID1A tiled peptide array. Predicted PLK1 consensus sites in TOX4-binding ARID1A peptides are shown**. B,** AlphaFold structural models of TOX4 with the two ARID1A peptides where TOX4 binding signal was detected. **C,** Co-immunoprecipitation of TOX4 and ARID1A. TOX4 was immunoprecipitated from RPE1 and HEK293T cells. IgG was used as a negative control. TOX4 immunoprecipitates were probed for the presence of ARID1A and ubiquitin. Short and long exposures are shown for ARID1A detection.

We used the AlphaFold protein–protein interaction prediction algorithm ^49^ to predict the structure of the ARID1A peptides complexed with TOX4. For both peptides we obtained low confidence output models with low interface-predicted template modelling (0.13 and 0.41 respectively) and high PAE scores, not unexpectedly, given the highly disordered nature of TOX4 and one of the ARID1A peptides (Figure 5B).

To validate the interaction, we immunoprecipitated TOX4 from RPE1 and HEK293T cells and probed the immunoprecipitates with an anti-ARID1A antibody (Figure 5C). We reproducibly detected anti-ARID1A reactive bands at 270 kDa band and above in TOX4 immunoprecipitates, confirming that TOX4 and ARID1A interact *in vivo*. Intriguingly, the ladder pattern of ARID1A bands was reminiscent of ubiquitination or sumoylation. We therefore probed TOX4 immunoprecipitates with an anti-ubiquitin antibody and observed that the ladder-like bands above 270 kDa were indeed ubiquitin-containing bands. These results confirm that TOX4 interacts with (poly)ubiquitinated ARID1A and suggest an interplay between (de)phosphorylation and ubiquitination of ARID1A.

### PRISMA uncovers ARID1A binding to CDK2/cyclin A2

The detection of CDK2 and cyclin A2 as overlapping consecutive binders in at least two different regions on ARID1A was of particular interest (Figure 6A). The highest binding signal for both proteins was detected on peptides spanning aas 1391-1430, where CDK2 and cyclin A2 binding overlapped on 2 out of 3 peptides. CDK2 also bound strongly between aas 531-600, with cyclin A2 displaying residual binding on one peptide in that span. Both binding sites overlapped with sequences of intrinsic disorder that were predicted to fold upon binding to a protein partner by the ANCHOR2 algorithm ^32^. CDK2 alone also displayed strong binding on peptides spanning aas 1451-1480, and residual binding on peptides spanning aas 331-370, which encompass a CDK2 consensus site. Phosphorylation of Serine 363 within this sequence is one of the most frequently detected ARID1A post-translational modifications (Figure S2). Exploration of a published CDK2-dependent phosphopeptidome study revealed that phosphorylation of Ser363 is dependent on CDK2 ^50^. Therefore, our results endorse the direct interaction between ARID1A with CDK2 and/or cyclin A via small regions of intrinsic disorder.

**Figure 6.**
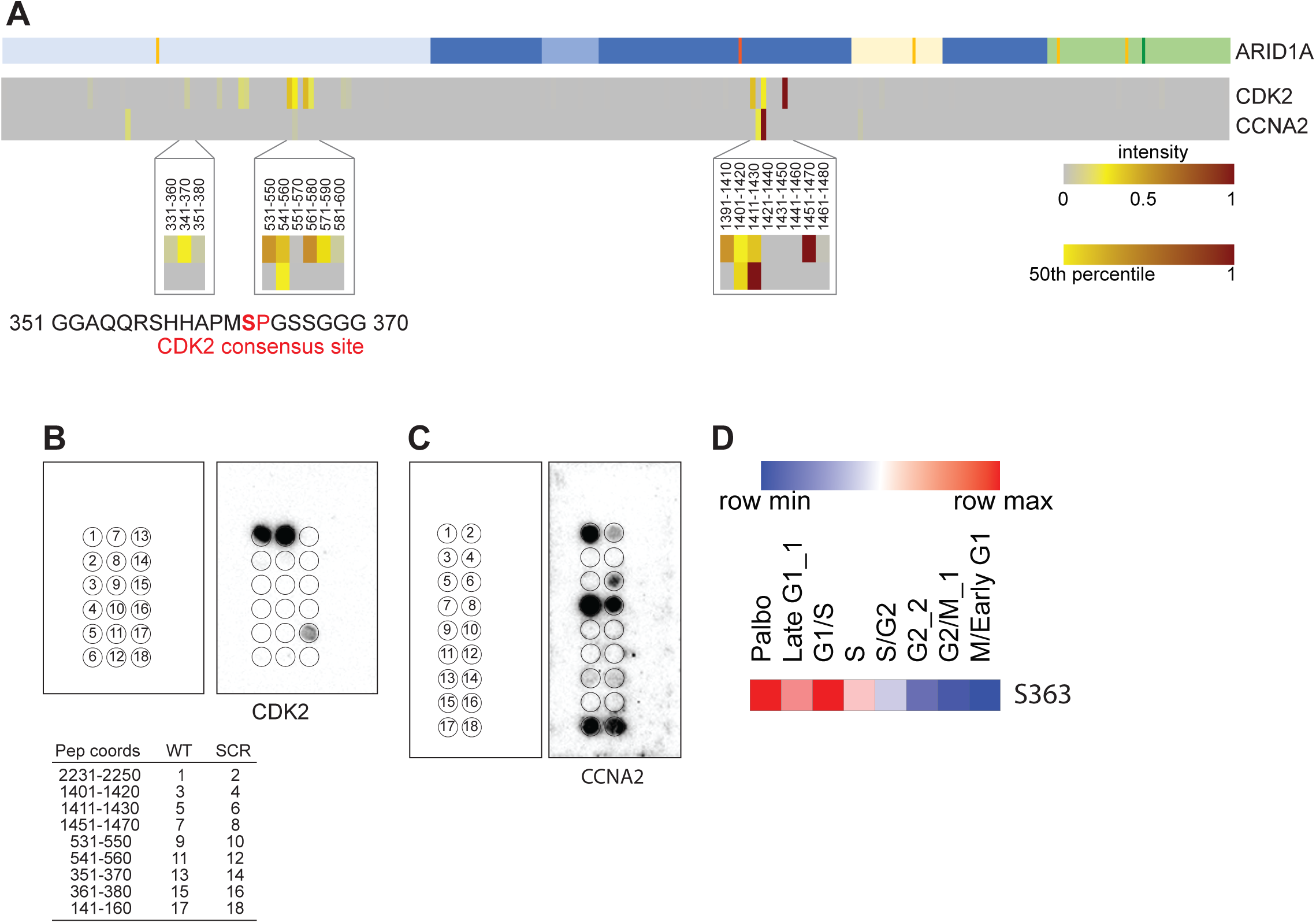
CDK2/Cyclin A bind to ARID1A. **A,** CDK2 and cyclin A2 PRISMA binding profiles with zoomed in sections of CDK2/CCNA2 binding overlap and CDK2 phosphorylation site on ARID1A. **B,** Validation of CDK2/CCNA2 binding to a subset of ARID1A peptides by far-western blotting using a recombinant mix of CDK2/CCNA2. WT, wild type peptide sequence; SCR, scrambled sequence control. **C,** Validation of CCNA2 direct binding to a subset of ARID1A peptides by far-western blotting using recombinant CCNA2. Peptides as in B. **D,** Heatmap showing levels of phosphorylated S363 (normalised to ARID1A abundance) along the cell cycle. See also Table S4.

To validate this, we synthesised a peptide subset array containing ARID1A peptides that had shown binding to CDK2 and/or CCNA2, an ARID1A peptide containing an RxL motif as positive control (RxL is the best characterised docking motif for cyclin A ^51^), and the corresponding scrambled sequences counterparts as negative controls (Table S4). The peptide array was then probed with a mix of recombinant full length CDK2 and a CCNA3 fragment spanning aas 174-432 that contains the hydrophobic patch that docks on the RxL motif ^51^ (Figure 6B). Western blot analysis revealed binding of CDK2 to the RxL peptide and not to its scrambled counterpart. We also detected binding of CDK2 to the ARID1A peptide spanning aas 1451-1470, but not to its scrambled counterpart. We could not perform cyclin A detection in this experiment because the fragment used does not contain the epitope of our anti-cyclin A antibody. To establish which of CDK2 and cyclin A is the direct binder to ARID1A, we probed a second copy of the array with recombinant full length CCNA2 (Figure 6C). Detection with an antibody against CCNA2 showed that CCNA2 alone binds to ARID1A peptide spanning aas 1451-1470, suggesting that CCNA2 mediates the binding of CDK2 to this sequence.

Interestingly, CDK2 or cyclin A2 have never been identified as ARID1A-associated proteins in our AP-MS experiments or any published ARID1A AP-MS studies ^4,52,53^. Thus, our data highlight the utility of the PRISMA strategy in revealing weak, transient and/or low abundance ARID1A interactions, possibly mediated by SLiMs, that are not detectable by traditional affinity purification strategies.

### Exploring the role of ARID1A Ser363 phosphorylation

A survey of our recent cell cycle phosphoproteomics study ^54^ revealed that ARID1A Ser363 phosphorylation is modulated along the cell cycle, being higher in G1 and S phases, and decreasing at the end of S phase and throughout G2 and mitosis (Figure 6D). To investigate the role of Ser363 phosphorylation, we generated RPE1 cells lines expressing inducible EmeraldGFP (emGFP)-tagged wild type (WT) ARID1A, phospho-mimetic Ser 363 to Asp mutated ARID1A (ARID1A_S363D) or phospho-dead Ser 363 to Ala ARID1A (ARID1A_S363A) in an ARID1A KO background. ARID1A_S363A was observed as a ∼250 kDa band in agreement with its predicted size, but the S363D mutant could not be detected with anti-ARID1A antibodies, and in fact was only expressed as a small protein fragment containing emGFP (Figure S4A, B). We confirmed this through primary sequence analysis by mass spectrometry of GFP-immunoprecipitates (Figure S4C). ARID1A peptides spanning the entire ARID1A sequence were detected in ARID1A_S363A immunoprecipitates. In contrast, in ARID1A_S363D immunoprecipitates we could only detect GFP peptides and ARID1A peptides upstream of S363D, indicating that the ARID1A_S363D protein detected by anti-GFP western blotting is a truncated form of ARID1A. This suggests that the ARID1A_S363D mutant protein is unstable and gets degraded to a truncated version in a rapid manner or was strongly selected against and a genomic rearrangement led to the truncated form. We could not reverse this degradation with proteasome inhibition (Figure S4B). Mass spectrometry analysis of GFP-trap immunoprecipitates also detected most BAF complex subunits in ARID1A_S363A immunoprecipitates (Figure S4D), suggesting mutation of Ser363 into Ala does not impact the ability of ARID1A to underpin the assembly of the complex. As expected, no other BAF subunits were detected in ARID1A_S363D immunoprecipitates, since the complex integrity and assembly relies on the ARID1A C-terminal domain ^55^. Whilst EmGFP-ARID1A_S363A fusion protein localised mostly to the cell nucleus (Figure S4E) in a similar manner to endogenous ARID1A, the S363D mutant localised to the cytoplasm, again as expected due to the loss of the NLS motif at position 1368-1387.

To test if S363 phosphorylation is important for cell growth, we induced the expression of emGFP, wild type and S363A mutant emGFP-ARID1A proteins in ARID1A KO RPE1 cells and monitored their growth by measuring cell confluency over 120 h (Figure 7A). We observed that in these conditions, cells expressing em-GFP-ARID1A_S363A had a mild proliferation defect compared to cells expressing wild type emGFP-ARID1A, suggesting that ARID1A S363 phosphorylation may be important for cell proliferation.

**Figure 7.**
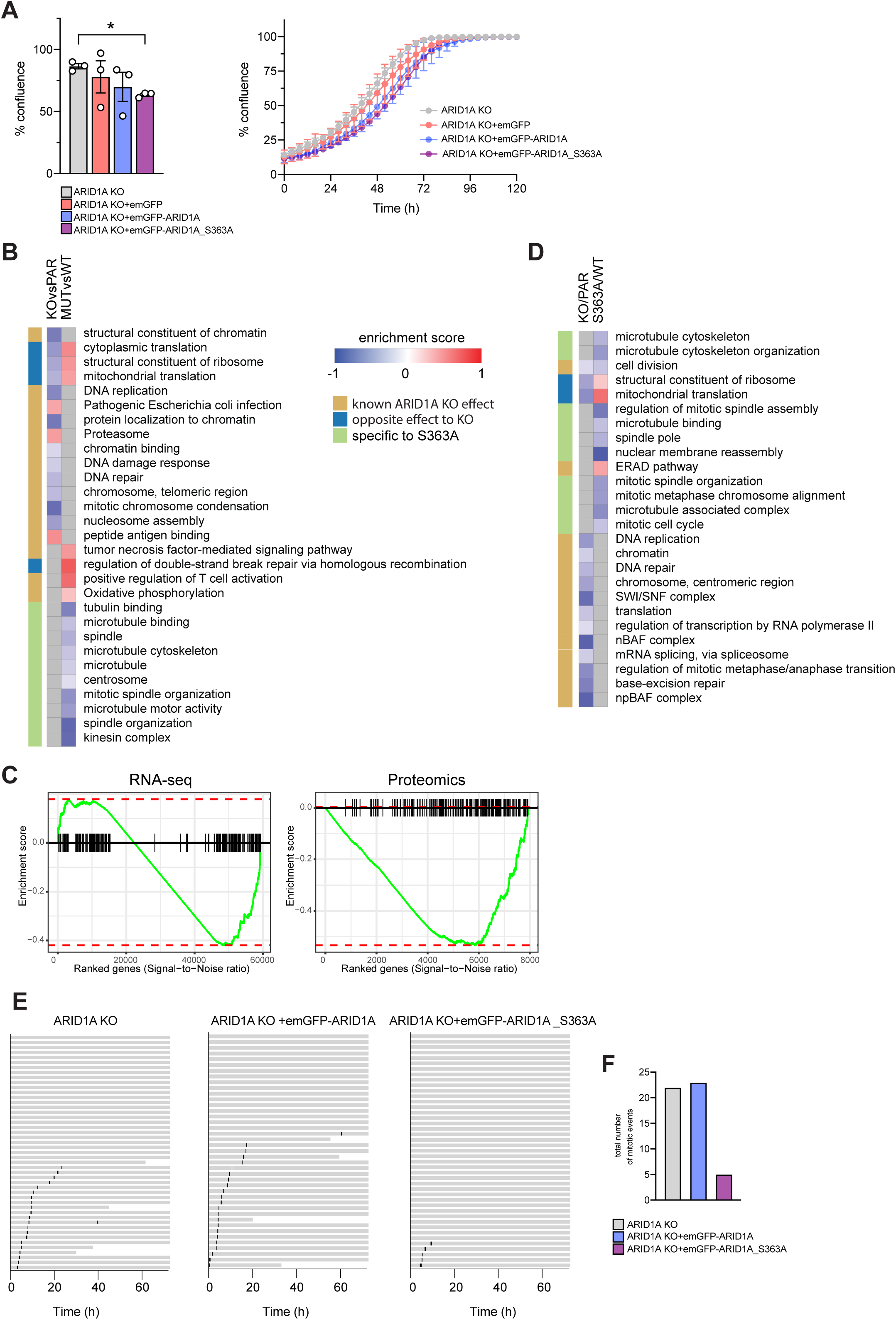
Mutation of ARID1A S363 to Alanine impacts ARID1A function. **A,** Bar graph and curve showing growth (as % confluence) of ARID1A KO cells alone or expressing emGFP only, wild type emGFP-ARID1A WT or emGFP-ARID1A_S363A. Points correspond to independent biological replicates, n = 3, mean ± SEM. **B,** Heatmap of enriched GO and KEGG terms from Gene Set Enrichment Analysis of RNASeq data from wild type ARID1A, ARID1A KO alone, and ARID1A KO expressing ARID1A wild type or S363A mutant form. **C,** Enrichment plot of genes (RNA-seq) and proteins (Proteomics) with the GO term Microtubule cytoskeleton annotation, showing lower levels of genes and proteins annotated with this term are associated with the ARID1A S363A mutation. **D,** Heatmap of enriched GO and KEGG terms from Gene Set Enrichment Analysis of proteomics data from wild type ARID1A, ARID1A KO alone, and ARID1A KO expressing ARID1A WT or S363A mutant form. **E,** Cell fate profiles comparing ARID1A KO, ARID1A KO expressing either emGFP-ARID1A WT or emGFP-ARID1A_S363A. Each track represents an individual cell, with distinct colours indicating specific cell cycle events. **F,** Bar graph showing mitotic events recorded over a 72-hour imaging period for ARID1A KO, ARID1A KO expressing either EmGFP-ARID1A WT or EmGFP-ARID1A S363A. See also Tables S5 and S6.

To explore the effects of the S363A mutation on ARID1A transcriptional regulation activity, we performed transcriptomics analysis using RNA sequencing (RNA-Seq) of ARID1A KO cells expressing emGFP-tagged wild type (WT) ARID1A, ARID1A_S363A or emGFP only, the untransfected ARID1A KO and the parental wild type cells after induction with doxycycline for 24 h. We then compared parental to ARID1A KO cells, and KO cells expressing wild type ARID1A to those expressing ARID1A_S363A. Differences between wild type and ARID1A-defficient cells were modest, as has previously been reported, and so were those between cells expressing induced wild type or mutant ARID1A. Gene set enrichment analysis (GSEA) revealed that ARID1A-deficient cells showed reduced expression of genes involved in chromatin organisation, replication, DNA repair, translation and peptide presentation compared to the parental (Figure 7B, Table S5), as has been reported previously ^11^. Notably, cells expressing ARID1A_S363A did not display these molecular phenotypes when compared to cells expressing WT ARID1A. They did however show increased expression of genes involved in oxidative phosphorylation, TNF signalling and interferon response, all of which are hallmarks of loss of ARID1A function ^56–59^. Strikingly, they also showed reduced expression of genes involved in microtubule-related processes, such as microtubule binding, mitotic spindle organisation, microtubule motor activity and kinesin complex (Figure 7B, C; Table S5).

To assess whether these effects translate on the protein level, we performed proteome profiling using quantitative MS. As expected, ARID1A KO cells displayed lower abundance of other BAF complex subunits at the protein level compared to the WT parental lines (Figure 7D) as previously shown ^12^, but this was not the case for EmGFP-ARID1A_S363A cells compared to cells expressing emGFP-ARID1A_WT, confirming that the S363A mutation does not affect the BAF complex assembly or stability. GSEA of proteomics data recapitulated the microtubule-based defects observed in the transcriptomics analysis (Figure 7C, D, Table S6). This is striking given that transcriptomic and proteomics changes elicited by loss of key SWI/SNF subunits are generally not well correlated ^11,12,60^. Cells expressing mutated ARID1A showed reduced abundance of proteins involved in microtubule organisation and mitotic events such as mitotic spindle assembly, chromosome alignment and nuclear membrane reassembly after mitosis. There was considerable overlap of downregulated genes/proteins annotated with “microtubule cytoskeleton” (50%) or “mitotic spindle organization” (75%) between RNA-seq and proteomics datasets. These results show that phosphorylation of S363 is important for specific aspects of ARID1A function related to the microtubule cytoskeleton.

The reduced abundance of proteins involved in microtubule dynamics in ARID1A_S363A mutant cells suggested that they may have defects in mitosis. To explore this, we induced expression of wild type or mutant ARID1A in ARID1A KO cells and monitored cell division using live-cell imaging over a period of 72 h (Figure 7E). We observed that cells expressing ARID1A_S363A rarely underwent mitosis over the 72 h, whilst cells expressing WT ARID1A divided at similar rates to the untransfected KO cells (Figure 7F). In summary, our results suggest that ARID1A Ser363 phosphorylation is important for normal mitosis and cell division.

## Discussion

Here we have used a peptide array interactomics strategy (PRISMA) to uncover novel interaction partners and binding sites of the SWI/SNF scaffold subunit ARID1A. This relatively new approach detects biophysical protein binding and has proven very useful to identify interactions that are mediated by SLiMs ^31,61–63^, but has only been applied to a small number of proteins to date. Our study significantly contributes to the understanding of molecular mechanisms of ARID1A regulation and function, by exposing novel ARID1A interacting partners and interaction sites, and providing a novel resource that can be used by the SWI/SNF community to further explore ARID1A biology.

Integration of SLiM annotations on the target protein with the PRISMA screen data can significantly contribute to the detection of confident interaction candidates for validation and follow-up experiments, enhancing the confidence in potential true interactions and providing insight into the regulation, molecular mechanism and function of protein interactions. Matching of ARID1A SLiMs to proteins containing the partnering globular domains highlighted SIN3A as a direct interactor through its PAH domain(s). Although ARID1A has been shown to be required for the recruitment of SIN3A to chromatin to regulate the expression of cell cycle and cytokine genes and *TERT* ^37,42,64^, and interactions of SIN3A with the BAF complex have been reported ^40,41,65^, only one study had previously demonstrated an interaction between SIN3A and ARID1A ^37^. Our data shows that the interaction between ARID1A and SIN3A is direct and mediated through the Sin3 SLiM.

The high concentration of peptide on the PRISMA arrays facilitates the detection of weak and transient interactions that are challenging to capture in traditional affinity purification approaches employing endogenous proteins. This may have been instrumental in the identification of novel interactor TOX4. The detection of the ubiquitin-containing ARID1A laddered pattern in the TOX4 immunoprecipitate suggests that TOX4 preferentially binds to (poly)ubiquitinated ARID1A *in vivo*, and this may explain why this interaction is not readily detected in traditional ARID1A IP-MS interactomes. Interestingly, the TOX4-binding ARID1A peptide encompassing aas 1691-1710 contains a predicted ABBA motif, a SLiM that has been shown to act as a degradation signal for key mitotic regulators and a signal for kinetochore recruitment ^66^, raising the question of whether the interaction with TOX4 plays a role in ARID1A degradation. PP1 regulatory subunit TOX4 has been shown to have a role in cell fate reprogramming and differentiation ^67,68^, and was reported to bind to DNA adducts generated by cisplatin ^69^. Seeing as it plays a role in fundamental processes including genome stability ^47,70^, it is no surprise that it has been shown to promote tumourigenesis and so has interesting therapeutic potential. Both PLK1 and GSK3 form reciprocal regulatory loops with PP1 ^71–73^, and our data suggests that the interplay of PP1 and either or both kinases might regulate some aspect of ARID1A function. Indeed, PLK1 inhibition was recently shown to be synthetic lethal with ARID1A deficiency ^74^.

Similarly, this seems to be the case with CDK2 and CCNA2, which have not been previously detected in any ARID1A interactome study. We detect weak binding of the kinase on a peptide containing a CDK2 consensus site that has previously been shown to be phosphorylated in a CDK2-dependent manner, and stronger binding, together with cyclin A, on peptides within the region of intrinsic disorder in ARID1A between the ARID and the C-terminal globular domains. The ARID1A peptide with the strongest CDK2/CCNA2 binding signal does not contain a canonical cyclin A docking motif (RxL). It is, however, reminiscent of a yeast cyclin binding motif PxxPxF ^75^. Intriguingly, there is an RxL motif in the C-terminal of ARID1A that overlaps with the SIN3 binding motif, that indeed binds cyclin A *in vitro* (Figure 6C).

Our exploration of published cell cycle phosphorylation data ^54^ suggests that ARID1A Ser363 is regulated in a cell cycle dependent manner and represents the first report of ARID1A regulation through phosphorylation. The mutation experiments prompted by findings from the PRISMA data indicate that the dynamic behaviour of Ser363 phosphorylation is specifically required for the expression of genes involved in microtubule-based processes, many of which are cell cycle genes. We find that expression of ARID1A_S363A mutant, where S363 cannot be phosphorylated, leads to decreased expression of genes and proteins involved in microtubule organisation and movement. Previous work has shown that ARID1A loss results in reduced expression of cell cycle and mitotic spindle genes at RNA and protein level ^37,76^, and our data suggests that ARID1A S363 dynamic phosphorylation specifically regulates this transcriptional program. Whilst SWI/SNF controls gene expression, many of the transcriptional changes observed when specific subunits are lost do not often correlate well with changes at the protein level ^11,12,60^. Importantly, our study shows that the regulation of MT-based genes by ARID1A S363 is cascaded down to the protein level and therefore results in a functional phenotypic effect on the cells. With ARID1A (and SWI/SNF) being master regulators of a huge number of genes and many different transcriptional programs, it is not surprising that regulation of these different sets of genes must happen through precise and specific mechanisms. This is not unprecedented for SWI/SNF. For instance, during mitosis SMARCE1 and SMARCB1 remain bound specifically to genes required for the completion of mitotic cell division and lineage-identity genes. By contrast, catalytic subunits SMARCA2 and SMARCA4 have been shown to localise to the nucleus through interphase but are displaced from condensed chromatin during mitosis ^77,78^. ARID1A has also been shown to be excluded from chromatin during mitosis ^78^. The genes annotated to microtubule processes that are affected by the S363A mutation are enriched in SIN3A targets (ENCODE ChIP data from EnrichR), and their expression along the cell cycle is positively correlated to that of CCNA2 and inversely to the phosphorylation of ARID1A S363 (data not shown). This raises the possibility that S363 (de)phosphorylation may regulate SIN3A chromatin recruitment and/or maintenance. Notably, ARID1A S363 phosphorylation is upregulated in breast cancer ^79^. Incidentally, this residue is in a stretch of sequence that is missing in ARID1B, an ARID1A paralog with an antagonistic role in cell cycle control ^37^.

Some well documented ARID1A interactions were not identified by our PRISMA assay. There are many reasons why this can happen, such as interactions being dependent on the presence of PTMs, cell line or context-dependent, or simply of too low abundance to be detected by PRISMA. It is also possible that missing interactions are reliant on globular structures on ARID1A that are not recapitulated by the short peptides used in our array. We also noted that PRISMA binding profiles of known ARID1A interacting proteins that are involved in DNA damage did not agree with published binding regions. This could reflect a dependency on DNA damage signalling, including PTMs on either ARID1A or interacting proteins.

Overall, our study has uncovered new biology on ARID1A, provides new avenues for further mechanistic exploration of ARID1A and SWI/SNF function and could inform therapeutic strategies impacting ARID1A function. Furthermore, our work represents a useful resource more broadly to investigate sequence or structural features governing protein interactions.

## Resource availability

Requests for further information and resources should be directed to and will be fulfilled by the lead contact, Mercedes Pardo. All unique/stable reagents generated in this study are available from the lead contact with a completed materials transfer agreement. RNA-sequencing datasets generated in this study are available in the GEO repository with accession number GSE313270. The mass spectrometry proteomics data have been deposited to the ProteomeXchange Consortium via the PRIDE ^80^ partner repository with dataset identifiers PXD074557 (chromatin profiling), PXD074744 (ARID1A_S363A proteome profiling) and PXDXXXXXX (ARID1A PRISMA).

## Acknowledgements

We thank the Institute of Cancer Research Flow Cytometry Facility, Insect Cell Culture Facility and Scientific Computing Team for support. J.S.C. and M.P. acknowledge funding from the Wellcome Trust (223745/Z/21/Z), BBSRC (BB/Y004477/1) and ICR core funding. J.A.D. acknowledges funding from Cancer Research UK DRCRPG-June24/100004. C.A., F.Y. and M.W. are supported by the Wellcome Career Development Award (302297/Z/23/Z).

## Author contributions

M.P. and J.C. conceived the project. M.P., C.M., K.A.L., F.S., L.S. and Z.K. performed experiments. M.P., C.M., and K.A.L. performed data analysis. M.W. and F.Y. contributed reagents. M.P., C.A., J.A.D. and J.C. obtained funding and provided input and supervision. M.P. prepared the original draft. M.P., C.M., K.A.L., C.A., J.A.D. and J.C. contributed to review and editing.

## STAR Methods

### Cell culture

hTERT-RPE1 (RPE1, ATCC) were cultured in Dulbecco Modified Minimal Essential Medium (DMEM)/F-12 (Merck) supplemented with 10% FBS (Gibco), 200µM glutamax (Gibco), 0.26% sodium bicarbonate (Gibco), and 1% penicillin/streptomycin (P/S)(Merck). HEK293TN (Systems Biosciences) were cultured in DMEM supplemented with 10% FBS and 1% P/S. All cells were maintained at 37°C in a humified incubator with 5% CO_2_ and were regularly tested for mycoplasma contamination.

### Plasmids

Emerald-GFP (emGFP) and emGFP-tagged ARID1A, ARID1A S363A and ARID1A C-terminus (aa. 1009-2285) were amplified by PCR with Q5® High-Fidelity DNA Polymerase (M0491S, NEB) using pLenti-puro-ARID1A plasmid (Addgene #39478) as template and cloned into pCW57-MCS1-2A-MCS2 plasmid (Addgene #71782) through NEBuilder® HiFi DNA Assembly (NEB, E2621S). Plasmids were prepared using the Qiagen Plasmid Plus kit as per the manufacturer’s protocol and stored at −20°C. Correct nucleotide sequence was confirmed by whole plasmid sequencing (Plasmidsaurus). The lentiviral packaging and envelope plasmids psPAX2 and pMD2.G were gifts from Professor Didier Trono (Addgene plasmids #12259 and #12260). Oligos used are listed in Table S8.

### Cell line generation

For lentivirus production, 10 µg of the target plasmid was transfected into HEK293TN cells along with the lentiviral packaging and envelope plasmids, psPAX2 and pMD.2G, using Lipofectamine 3000 (Thermo) according to the manufacturer’s protocol. Viral particle-containing medium was harvested at 48 hr and 72 hr post transfection, combined, centrifuged for 5 min at 300 *x g* to remove debris, and filtered through 0.45 µm cellulose acetate filters (Sartorius). The resulting supernatant was concentrated using the Lenti-X Concentrator (Takara Bio) according to the manufacturer’s protocol. For transduction, concentrated viral supernatant was added to RPE1 cells in 6-well plates along with 6 µg/mL polybrene (Merck). Cells were expanded to 10 cm^2^ dishes, and 1 µg/mL DOX was added to cells to induce protein expression. After 24 hr, the pool of GFP-positive cells was sorted into 6-well plates by the BD FACSAria III sorter (BD Biosciences). Protein expression in the resulting cell lines was confirmed by Western blot analysis and immunofluorescence microscopy.

### Compounds

Doxycycline hyclate (DOX, Merck) was dissolved in RNase-free H_2_O to a stock concentration of 1 mg/mL. 5-ethynyl-2’-deoxyuridine (EdU, Invitrogen) was dissolved in DMSO (Merck) to a stock concentration of 10 mM.

### Peptide arrays

Cellulose acetate membranes containing immobilised PRISMA peptides spanning the whole length of ARID1A were purchased from JPT Peptide Technologies (Berlin, Germany). Peptide sequences are listed in Table S1.

Peptide arrays for the CDK2/CCNA2 validation experiments (Table S2) were prepared in-house. Peptides were synthesised on derivatised cellulose membranes using a Multipep1 instrument (CEM Microwave Technology Ltd.) by cycles of coupling of N(a)-Fmoc-protected aminoacids (Sigma-Aldrich) through carboxylic group activation with diisopropylcarbodiimide (Sigma-Aldrich) and Oxyma pure (CEM), followed by deprotection of alpha-amino groups by piperidine (Sigma-Aldrich), followed by acetylation of N-terminal residues with acetic anhydride (Sigma-Aldrich). All peptides were N-terminally acetylated. After peptide synthesis, side chain deprotection was performed as previously described ^81^.

### Full length ARID1A PRISMA

The PRISMA screen was carried out a single biological replicate with technical replicate MS measurements. Nuclear isolation was performed by lysing wild type RPE1 cells in an isotonic buffer (10 mM Tris-HCl, 10 mM NaCl, 1.5 mM MgCl2, 0.34 M sucrose containing 1 mM dithiothreitol (DTT) and Halt Protease inhibitors (Thermo) and 0.05% NP-40 as described previously ^82^. Nuclei were lysed by incubation for 10 mins on ice with PRISMA high salt lysis buffer (50 mM HEPES pH 8.0, 450 mM NaCl, 1 mM EDTA, 10% glycerol, 0.5% NP-40, 1 mM DTT, Halt protease inhibitors) followed by Dounce homogenisation. The nuclear lysate was incubated with universal nuclease (Pierce) for 1 hour on ice and then centrifuged at 13000 rpm for 15 min at 4°C. The cleared nuclear lysate was diluted 1:3 with PRISMA dilution buffer (50 mM HEPES pH 8.0, 1 mM EDTA, 10% glycerol, 0.5% NP-40, 0.3 mM DTT, Halt Protease inhibitors (Thermo). The PRISMA membrane was wetted for 15 mins in PRISMA detergent-free wash buffer (50 mM HEPES pH 8.0, 150 mM NaCl, 1 mM EDTA, 10% glycerol), followed by blocking with yeast tRNA at 1 mg/ml (Life Technologies) in the same buffer. The membrane was then washed with PRISMA low salt lysis buffer (50 mM HEPES pH 8.0, 150 mM NaCl, 1 mM EDTA, 10% glycerol, 0.5% NP-40) 5 times for 5 mins. The membrane was then incubated with 40 mg of RPE1 nuclear extract at ∼1.4 mg/ml in the diluted lysis buffer for 1h at 4°C. After removal of the nuclear lysate, the membrane was first washed with PRISMA low salt lysis buffer 3 times for 5 mins, followed by PRISMA detergent-free wash buffer 3 times for 5 mins, and then air dried for 1h. The peptide spots were excised and placed one to a well in a 96-well HTS filter plate (Millipore) containing 200 μl of 5 mM TCEP (Sigma-Aldrich) in 50 mM ammonium bicarbonate. Excised membrane spots were incubated for 15 mins at 37°C with shaking to allow reduction, followed by alkylation with 10 mM iodoacetamide for 30 mins shaking at room temperature in the dark. TCEP and iodoacetamide were removed by spinning the plate at 1000 rpm for 1 min, and spots were washed 3 times with 50 mM ammonium bicarbonate. Samples were digested in the filter plate with 0.5 μg of trypsin (sequencing grade, Roche) and 0.25 μg of LysC (Wako Chemicals) per spot/well for 2 hours at 37°C, followed by overnight incubation at room temperature. Peptides were collected by centrifugation onto a fresh plate, and the filter plate stack was flushed twice with 50 μl of 100% HPLC methanol and the flowthrough pooled with the first peptide collection. The peptides were dried and then resuspended in 0.5% formic acid ready for MS analysis.

### PRISMA mass spectrometry analysis

Peptides were analysed with online nanoLC-MS/MS on an Orbitrap Fusion Tribrid mass spectrometer coupled with an Ultimate 3000 RSLCnano System. Samples were first loaded and desalted on a nanotrap (100 µm id x 2 cm) (PepMap C18, 5 µm, 100A) at 10 µl/min with 0.1% formic acid for 10 min and then separated on an analytical column (75 µm id x 50 cm) (PepMap C18, 5 µm, 100A) over a 60 min linear gradient of 4-32% acetonitrile (ACN):0.1% formic acid at 300 nL/min. This was followed by a 5 min column wash with 85% ACN:0.1% formic acid on the analytical column and a 10 min trap-column wash of a cleaning mixture containing 25% ACN:25% Isopropanol:25% methanol: 25% water. The total cycle time was 90 min.

The Orbitrap Fusion was operated in standard data-dependent acquisition. For peptide analysis, the precursors were fragmented in HCD (higher collision dissociation) cell at 32% collision energy with a quadrupole isolation width of 0.7 Th (Thomson unit), detected in the ion trap at a rapid scan rate, AGC of 4 × 10e5, and maximum injection time of 50 ms. Targeted precursors were dynamically excluded for further isolation and activation for 45 secs with 7 ppm mass tolerance. Peptide fragments were detected in the ion trap with a rapid scan rate setting. Technical replicate MS measurements were acquired for 50% of peptide spots.

### PRISMA-MS data analysis

The PRISMA raw MS data were analysed with MaxQuant software v. 2.0.3.0 in one batch and searched against the human Uniprot reference database (v. October 2022). A maximum of 2 missed cleavages were allowed, variable modifications were set to methionine oxidation, pyro-glutamic acid conversion of asparagine and glutamine, and N-terminal protein acetylation, and cysteine carbamidomethylation was set as fixed modification. The label free quantification (LFQ) and match between runs options were activated.

Further data processing was carried out using Perseus and Excel. Technical replicate measurements were integrated by averaging the LFQ intensity where two values were available, and the single value was used if only one was measured. Only proteins that were identified by two or more peptides were taken forward. Intensity values for the 228 peptides in each of the data rows were filtered according to the outlier filtering criterion, where the values were replaced by 0 if they were smaller than the 75^th^ percentile of the intensity value distribution of the protein ^31^. For visualisation and clustering, each data row was normalised between 0 and 1. Proteins that showed a consecutive binding pattern across a minimum of two matrix peptides were marked as making up the “core interaction dataset”.

Peptide properties were calculated using the Peptides R package v.2.4.4 ^83^. Annotation of GO terms was carried out with Perseus v.2.0.7.0 using an annotation file generated from Uniprot in October 2022. GO term and protein domain enrichment was carried out using the DAVID functional annotation tool ^84,85^. Venn diagrams were created with Venny 2.1 (bioinfogp.cnb.csic.es/tools/venny/).

### Preparation of cell lysates

For western blotting, cell pellets were resuspended in the appropriate volume of lysis buffer (10% glycerol, 50mM Tris-HCl (pH 7.4), 0.5% NP-40, 150mM NaCl) containing 0.25U/µL benzonase nuclease (Merck), 1x cOmplete™ EDTA-free protease inhibitor cocktail (Roche), and 1x PhosSTOP™ phosphatase inhibitor cocktail (Roche). Cells were lysed on ice for 45 minutes followed by centrifugation for 15 minutes at 16,000 *x g* at 4°C. The resulting supernatant containing whole cell protein extracts was collected.

Insect cell pellets were lysed in 50 mM Tris-HCl pH 8, 150 mM NaCl, 0.1% NP40, 1mM EDTA, 1x Halt protease and phosphatase inhibitor (Pierce) and 1 mM DTT on ice for 10 min, homogenised using a Dounce homogeniser and centrifugation for 15 minutes at 16,000 *x g* at 4°C.

For TOX4 immunoprecipitation, cell lysates were prepared as previously described ^86^.

### Immunoprecipitation

Relevant antibodies and non-relevant IgG control (normal)antibodies (1–10 μg) were coupled to 25-50 µl of Protein G Dynabeads (Life Technologies) following manufacturer’s protocols. Antibodies are listed in Table S7. Antibody-coupled beads were incubated with whole cell or nuclear lysates for 1–2 h at 4 °C, lysates were removed, and beads were washed with IPP150 buffer (150 mM NaCl, 10 mM Tris-HCl, 0.1% NP-40). Immunoprecipitated proteins were eluted by incubating the beads in 1x LDS sample loading buffer (Life Technologies) containing 50 mM DTT at 70 °C for 10 min. The cell lysate (input) and eluted samples were analysed by western blotting.

### Western blotting and Far Western on peptide arrays

Protein samples were separated in NuPAGE 4-12% BisTris or 3-8% Tris Acetate gels using MOPS or Tris Acetate running buffer (Life Technologies) respectively and transferred to nitrocellulose membranes (Cytiva). Blocking was performed with 5% milk in Tris Buffered Saline-0.1% Tween (TTBS) for 1 h or 10% milk in TTBS for 10 mins. Membranes were incubated in the appropriate primary antibody diluted in 5% milk blocking buffer at 4°C overnight, washed 3 times with TTBS buffer, and incubated in the appropriate horseradish peroxidase (HRP)-conjugated secondary antibody diluted in blocking buffer for 1 hour at room temperature. Blots were developed with ECL Prime, ECL Ultra (Cytiva) or Immobilon Forte (Merck) and chemiluminescence signal acquired on an iBright CL750 imager (Life Technologies).

For far western experiments, membranes containing peptide spots were first incubated in TTBS for 15 min, then blocked with 5% milk in TTBS for 1 h. Cyclin A (Bio-techne) at 5 ug/ml or CDK2/CCNA3[174-432] at 10 ug/ml in blocking solution were added to the membrane and incubated for 4 h at room temperature. Primary and secondary antibody incubations were performed at the corresponding dilutions in 5% milk in TTBS. Blots were developed with ECL Prime (Cytiva) and chemiluminescence signal acquired on an iBright CL750 imager (Life Technologies). Antibodies are listed in Table S7.

### Expression of proteins in insect cells

Codon-optimised human SIN3A (residues 449-1086) fused to StrepII tag and ARID1A-V5 (from Addgene plasmid #39311) cDNAs for insect cell expression were synthesised and subcloned into the pFastBac1 vector (GenScript). The SIN3A and ARID1A constructs were transformed into DH10MultiBac cells for bacmid generation, and bacmid DNAs were isolated by isopropanol precipitation. Bacmid DNAs were transfected into Sf9 (*Spodoptera frugiperda*) cells using CellFectinII (Thermo), and baculovirus stocks were amplified through two consecutive rounds of infection in Sf9 cells. For protein expression, Sf9 cells at a density of 1.5 × 10e6 cells/mL in Sf-900 III medium supplemented with penicillin/streptomycin were infected with the amplified virus at 1:30 (v/v). After 72 h incubation at 27°C with shaking at 130 RPM, cells were harvested and flash-frozen in liquid nitrogen for long term storage at −80°C until processing.

### Purification of CDK2 and cyclin A3

Both CDK2 (full length) and cyclin A3 (residues 174–432) were induced and expressed individually with 1 mM IPTG in BL21(DE3) Star cells at 18 °C overnight. Pellets containing GST-tagged CDK2 and His-tagged cyclin A3 were combined (two litres of Cdk2 cell culture and 2 litres of cyclin A3 cell culture) and resuspended in lysis buffer (50 mM Hepes pH8.0, 150 mM NaCl, 2 mM MgCl_2_, 5 % glycerol, 0.5 mM TCEP) supplemented with 5 units per ml of benzonase and cOmplete™ EDTA-free protease inhibitors (Roche). After sonication of the cell suspension, the cell lysate was centrifuged at 21,000 rpm. for 1 h at 4 °C and the supernatant was loaded onto pre-equilibrated 5 ml GSTrap™ HP column (Merck). The column was washed with lysis buffer and then the proteins were eluted out with elution buffer (20 mM reduced glutathione added into the lysis buffer). The eluted proteins were buffer exchanged into lysis buffer to remove reduced glutathione and GST-tag of CDK2 was then cleaved off with 3C PreScission protease overnight at 4 °C. After 3C protease cleavage, the sample was loaded onto 5 ml GSTrap™ HP column again, the flow-through was collected, concentrated and loaded onto pre-equilibrated size exclusion column (Superdex75 16/60 column, GE Healthcare). The protein complex was eluted out with gel filtration buffer (20 mM Hepes pH 8.0, 150 mM NaCl, 0.5 mM TCEP) and concentrated to final concentration 1.4 mg/ml. The purified protein was flash frozen and stored in −80 °C.

### Chromatin extraction

Cell pellets were resuspended in isotonic nuclear extraction buffer for chromatin extraction (15 mM Tris-HCL pH 7.5, 60 mM KCl, 15 mM NaCl, 5 mM MgCl2, 1 mM CaCl2, 250 mM Sucrose, 1 mM DTT, Halt Protease inhibitors) and 0.3% NP-40 was added dropwise to final concertation of 0.03% (Sigma-Aldrich), mixed gently and incubated on ice for 5 mins. Nuclei were pelleted at 400-600 rcf for 5 mins at 4°C. Nuclei were washed again with nuclear isolation buffer for chromatin extractions. Small fraction was verified to contain nuclei by light microscopy. Isolated nuclei were resuspended in hypotonic chromatin isolation buffer (3 mM EDTA, 0.2 mM ethylene glycol tetraacetic acid, 1 mM DTT, Halt Protease inhibitors (Thermo) and incubated on ice for 1 h, vortexing every 5-10 mins. Insoluble chromatin fraction was collected by centrifugation for 4 min at 1,700 rcf. Fraction was washed 2 times in hypotonic chromatin isolation buffer. Chromatin fractions were frozen on dry ice and stored at −20°C. Samples were re-suspended in the chromatin preparation buffer (100 mM Triethylamonium bicarbonate (TEAB) (Sigma-Aldrich), 1% sodium deoxycholate (SDC), 10% isopropanol, 50 mM NaCl, Halt Protease and Phosphatase inhibitor (Thermo) supplemented with and Benzonaze (Pierce). The chromatin pellet was disrupted with brief probe sonication and the samples were incubated on ice for 30 min. 30 μg of each sample was labelled TMTpro™ 16plex Label Reagent Set (Thermo) according to manufactures instructions.

### Chromatin profiling mass spectrometry

Offline peptide fractionation was based on high pH reverse-phase (RP) chromatography using the Waters XBridge C18 column (2.1 mm × 150 mm, 3.5 μm) on a Dionex UltiMate 3000 HPLC system at a flow rate of 0.2 ml/min. Mobile phase A was 0.1% (v/v) ammonium hydroxide, and mobile phase B was acetonitrile, 0.1% (v/v) ammonium hydroxide. Pooled TMT-peptides were resuspended in 200 μL of buffer A, centrifuged at 14 000 rpm for 5 min, and the supernatant was injected for fractionation with the following gradient: isocratic for 5 min at 5% phase B, gradient for 40 min to 35% phase B, gradient to 80% phase B in 5 min, isocratic for 5 min, and re-equilibrated to 5% phase B. The peptides were then resuspended with 0.1% formic acid and finally pooled into four fractions by combining the first and the last fractions.

LC-MS analysis was performed on the Dionex UltiMate 3000 UHPLC system coupled with the LTQ Orbitrap Lumos mass spectrometer (Thermo). Samples were analysed with the EASY-Spray C18 capillary column (75 μm × 50 cm, 2 μm) at 50 °C. Mobile phase A was 0.1% formic acid and mobile phase B was 80% acetonitrile, 0.1% formic acid. The gradient separation method was as follows: 150 min gradient up to 38% B, for 10 min up to 95% B, for 5 min isocratic at 95% B, re-equilibration to 5% B in 10 min, for 10 min isocratic at 5% B.

MS1 scans were acquired at a resolution of 120,000 over an m/z range of 375–1500 in the Orbitrap, with an AGC target of 4 × 10⁵ and a maximum injection time of 50 ms. Precursors were selected using Top Speed mode with a 3 s cycle time and isolated with a quadrupole isolation width of 0.7 Th. Collision energy for fragmentation via collision induced dissociation (CID) was set at 35%, with AGC target of 1 × 10⁴ and maximum injection time of 50 ms. MS3 spectra for TMT-based quantitation were acquired using synchronous precursor selection (SPS) of the top seven CID fragment ions. SPS-selected ions were fragmented by HCD with a collision energy of 35%. MS3 spectra were acquired in the Orbitrap over an m/z range of 100–500 at a resolution of 45,000, with an AGC target of 1 × 10⁵ and maximum injection time of 105 ms. Dynamic exclusion was activated for 45 s with a mass tolerance of 7 ppm.

### Chromatin profiling data analysis

MS raw data from IPs was analysed using Proteome Discoverer 2.4 (Thermo). Sequest search engine was used for protein identification and quantification followed by Percolator. Peptides were filtered at q-value <1% based on a decoy database search. The searches were performed against the full UniProt Human database (reference entries, January 2022) together with the cRAP contaminant database (ftp://ftp.thegpm.org/fasta/cRAP). Precursor mass tolerance was set to 20 ppm. Fragment mass tolerance was set to 0.5 Da. TMTpro (K and peptide N-terminus) and carbamidomethylation (C) were set as static modifications, while oxidation (M) and deamidation (NQ) were set as dynamic modifications. Protein abundances were normalised by multiplying each value by the maximum median of abundances across all channels and then dividing by the median of abundances of the corresponding channel. The mass spectrometry proteomics data have been deposited to the ProteomeXchange Consortium via the PRIDE ^87^ partner repository with the dataset identifier PXD074557.

### Incucyte proliferation assay and cell fate profiling

Cells were seeded in triplicate into 96-well plates at a density of 1,000 cells/well, and allowed to attach and grow for 24 hours. When indicated, 1 µg/mL DOX was then added to cells, and plates were transferred to the Incucycte SX5 (Sartorius) maintained at 37°C in a humidified 5% CO2 atmosphere. For tracking cell growth, phase contrast and fluorescence images (4 images per well) were collected every 4 hours up to a total of 120 hours. The average of three technical replicates was used, and % confluence (% of area covered by cells) was used as a readout of proliferation. For cell fate profiling, phase contrast and fluorescence images (4 images per well) were collected every 10 minutes up to a total of 72 hours. Image sequences were exported, and individual cells were analysed manually to generate cell fate profiles.

### Sulforhodamine B (SRB) proliferation assay

Cells were seeded in triplicate into 96-well plates at a density of 1,000 cells/well, and allowed to attach and grow for 24 hours. When indicated, 1 µg/mL DOX was then added to cells and cells were left to incubate for 6 days in total. After this, cells were fixed by the addition of 100 µl 10% TCA (Merck) and incubated at 4°C for at least 24 hours. Plates were then washed five times with water and allowed to completely dry before staining with 0.57% SRB powder (Merck) dissolved in 1% acetic acid and incubated for 1 hour room temperature. Unbound dye was removed by washing with 1% acetic acid, and the plate was allowed to dry completely. Protein-bound dye was solubilised in 10 mM Tris-HCl pH 9.5 for 30 minutes at room temperature with gentle rocking. Absorbance at 565 nm was quantified using the SpectraMax Microplate Absorbance Reader (PerkinElmer). The average of three technical replicates was used, and proliferation was defined as a percentage of the absorbance measured in treated wells compared with control wells.

### RNA-sequencing

For RNA-sequencing, cells were treated with 1 µg/mL DOX for 24 hours before harvesting. Pellets were harvested by scraping cells in ice-cold PBS following by centrifugation at 7,500 *x g* for 10 minutes. Three independent biological replicates were harvested for all cell lines. Pellets were resuspended in 500 µL TRIzol reagent and total RNA was extracted using a Direct-zol RNA miniprep kit (Zymo Research) according to the manufacturer’s protocol. RNA concentration and quality was confirmed with the High Sensitivity RNA ScreenTape (Agilent), using the TapeStation 4150 System (Agilent). Library preparation and sequencing was performed by Novogene Corporation Ltd. Novogene NGS Stranded RNA Library Prep Set was used to generate 250-300 bp insert strand specific libraries, and ribosomal RNA was removed using TruSeq Stranded Total RNA Library Prep. 50 million 150 bp paired-end reads were sequenced on an Illumina NovaSeq 6000.

### RNA-sequencing analysis

Fastq reads were checked using FastQC (v0.11.9) ^88^ and trimmed using TrimGalore (v0.6.6) ^89^. Residual ribosomal RNA reads were removed using Ribodetector with -e norRNA setting (v0.2.7) and strandedness was detected using RSeQC (v4.0.0). Reads were aligned to the human hg38.14 genome (GenBank GCA_000001405.29) using STAR alignment software (v2.7.6a) and reads mapping to genes were quantified using HTSeqCount (v0.12.4). Differential analysis of gene expression was calculated in R using *DESeq2* (v1.38.3).

### Whole cell proteomic profiling

Cell pellets were lysed in buffer containing 1% (w/v) sodium deoxycholate (SDC), 100 mM triethylammonium bicarbonate (TEAB), 10% (v/v) isopropanol, and 50 mM NaCl, freshly supplemented with 5 mM TCEP (Thermo, Bond-Breaker), 10 mM iodoacetamide (IAA), universal nuclease (1:2000, v/v; Pierce, #88700), and Halt protease and phosphatase inhibitor cocktail (1×; Thermo, #78442). Lysis was assisted by 5 min of bath sonication. Protein concentration was determined using the Quick Start Bradford protein assay (Bio-Rad). Aliquots containing 30 µg of total protein were digested overnight at room temperature with trypsin (Pierce) at a 1:20 (enzyme:substrate) ratio. Peptides were labelled with TMTpro reagents (Thermo) by adding 5 µL of reagent to 15 µL of peptide solution. Following labelling, samples were acidified to a final concentration of 2% formic acid, and precipitated SDC was removed by centrifugation.

The combined TMT-labelled peptide mixture was fractionated by high-pH reversed-phase chromatography using an XBridge C18 column (2.1 × 150 mm, 3.5 µm; Waters) on an UltiMate 3000 HPLC system over a 35-min gradient at 1% slope. Mobile phase A consisted of 0.1% (v/v) ammonium hydroxide, and mobile phase B consisted of 0.1% (v/v) ammonium hydroxide in acetonitrile. LC–MS analysis was performed on a Vanquish Neo UHPLC system coupled to an Orbitrap Ascend mass spectrometer (Thermo). Peptides were separated on a 25 cm capillary column (nanoEase MZ PST BEH130 C18, 1.7 µm, 75 µm × 250 mm; Waters) using a 110-min gradient from 5–35% mobile phase B (80% acetonitrile, 0.1% formic acid). Samples were first loaded onto a PepMap 100 C18 trapping column (5 µm, 0.3 × 5 mm, 1500 bar) before separation on the analytical column, which was coupled to a stainless-steel emitter via a Nanospray Flex ion source.

MS1 spectra were acquired in the Orbitrap at a resolution of 120,000. Precursors were selected for HCD fragmentation in top-speed mode (3 s cycle time) with a normalized collision energy of 32% and detected in the ion trap using turbo scan rate. MS3 spectra were triggered by real-time search (RTS) against a FASTA database containing reviewed UniProt *Homo sapiens* canonical and isoform sequences, using multi-notch isolation (10 notches) and HCD fragmentation at 65% normalized collision energy, with detection in the Orbitrap at 90,000 resolution. Dynamic exclusion was enabled for 45 s with a mass tolerance of 10 ppm, RTS close-out was set to a maximum of four peptides per protein, and targeted precursors were excluded from further activation. Static modifications used for RTS included TMTpro16plex on peptide N-termini and lysine residues and carbamidomethylation of cysteine. Variable modifications included oxidation of methionine and deamidation of asparagine and glutamine, with a maximum of one missed cleavage and up to two variable modifications per peptide.

Raw data were searched using the Sequest HT and Comet nodes implemented in Proteome Discoverer version 3.0 (Thermo) against a reviewed UniProt Homo sapiens FASTA database. Precursor mass tolerance was set to 20 ppm, and fragment ion mass tolerance was set to 0.5 Da for Sequest HT and 1 Da for Comet, allowing up to two missed tryptic cleavages. TMTpro modifications on peptide N-termini and lysine residues, as well as carbamidomethylation of cysteine, were specified as static modifications, while oxidation of methionine and deamidation of asparagine/glutamine were specified as dynamic modifications. Peptide-spectrum matches were filtered using Percolator with a peptide-level false discovery rate (FDR) of 1% based on a target–decoy strategy. Only unique peptides were used for protein quantification, considering protein groups for peptide uniqueness, and peptides with an average reporter ion signal-to-noise ratio greater than 3 were included in quantitative analysis. Data are available via ProteomeXchange with identifier PXD074744.

### Bioinformatics analysis of transcriptomics and proteomics data

Peptide properties were calculated using the Peptides R package v.2.4.4 ^83^. GO term and protein domain enrichment analysis of the ARID1A PRISMA core dataset was carried out using the DAVID functional annotation tool ^85^. Gene set enrichment analyses for whole cell proteomic profiling and RNA-seq data were performed using the 1D annotation enrichment algorithm ^90^ within the Perseus software platform ^91^. GSEA plots were generated in R using the *fgsea* package ^92^.

