## Supplementary material for "High resolution interaction surface mapping by PRISMA reveals novel ARID1A interactions": Figure S1

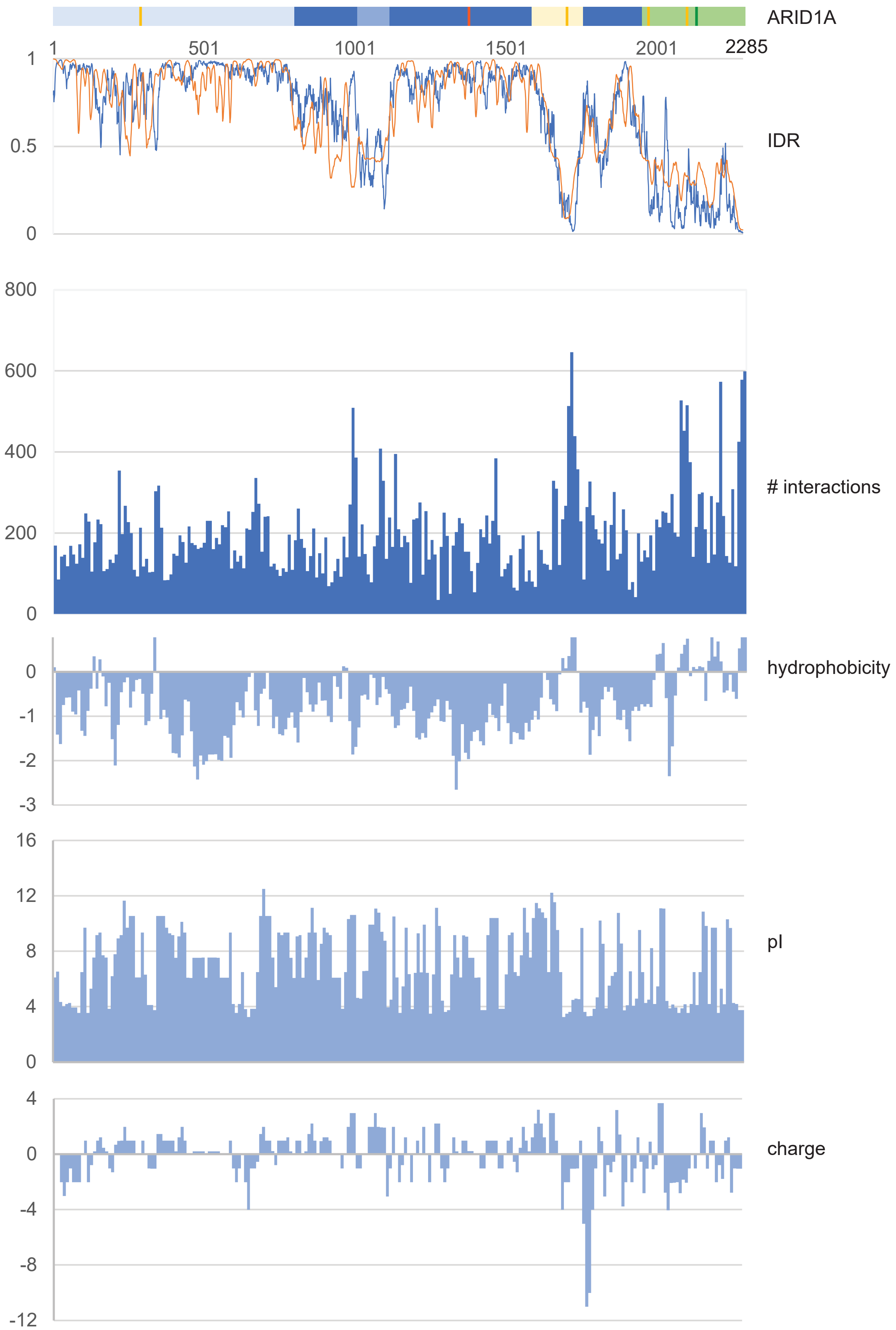


**Figure S1.** ARID1A PRISMA core interaction set profile depicting number of binding proteins identified per peptide, and graphs representing peptide properties, aligned to ARID1A domain diagram and intrinsic disorder region (IDR) prediction plot from IUPred2A. ARID1A domains and intrinsic disorder plots are as in Figure 1.
