## Supplementary material for "High resolution interaction surface mapping by PRISMA reveals novel ARID1A interactions": Figure S2

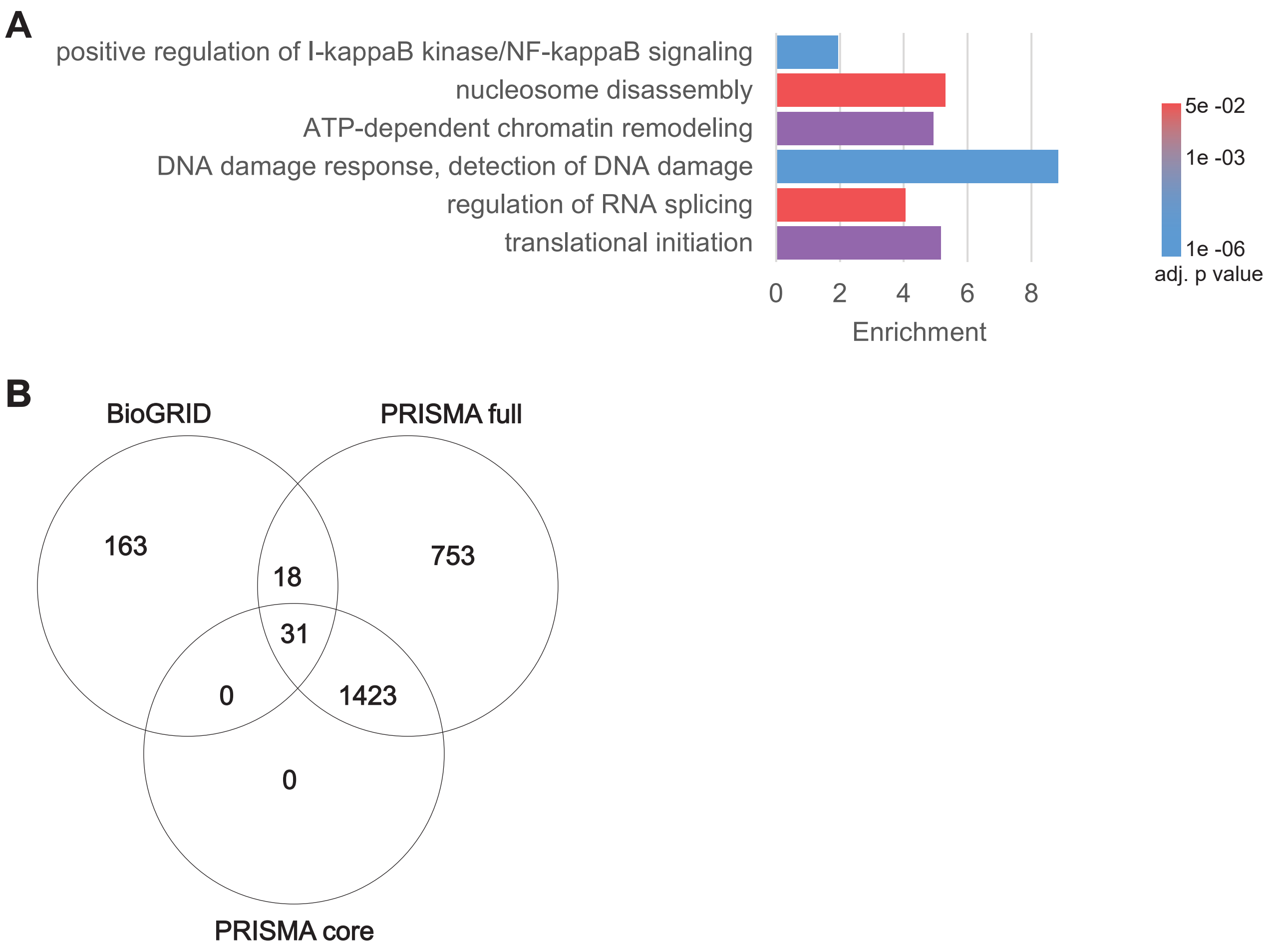


**Figure S2. A**, GO term enrichments of the PRISMA core interaction dataset. **B**, Venn diagram of the overlap between the ARID1A physical interactome (from BioGRID) and the PRISMA interaction set.
