## Supplementary material for "High resolution interaction surface mapping by PRISMA reveals novel ARID1A interactions": Figure S3

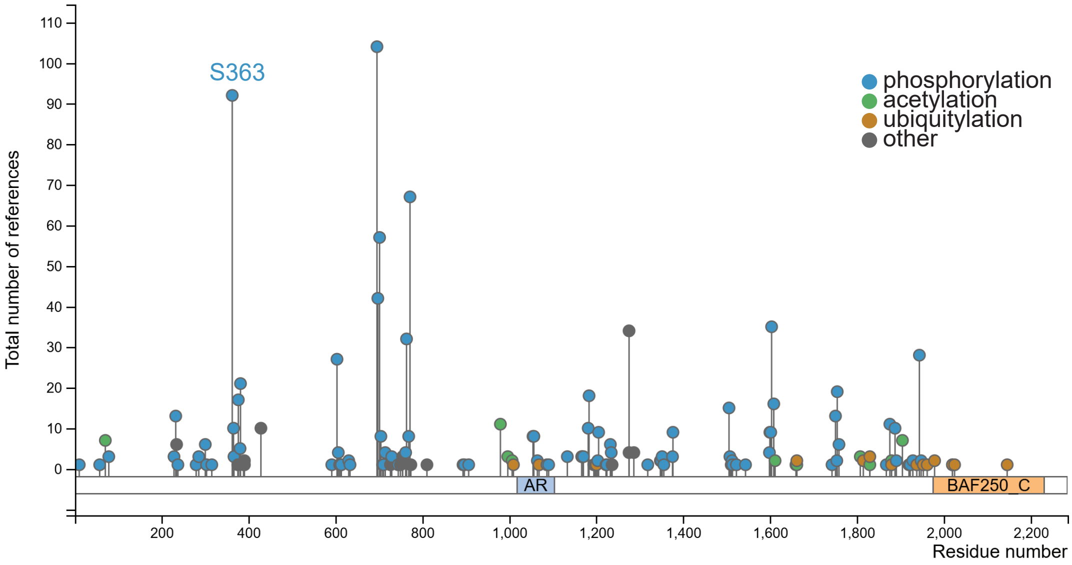


**Figure S3.** Graph showing all ARID1A post-translational modifications recorded in PhosphositePlus.org (to 01.10.2025).
