## Supplementary material for "High resolution interaction surface mapping by PRISMA reveals novel ARID1A interactions": Figure S4

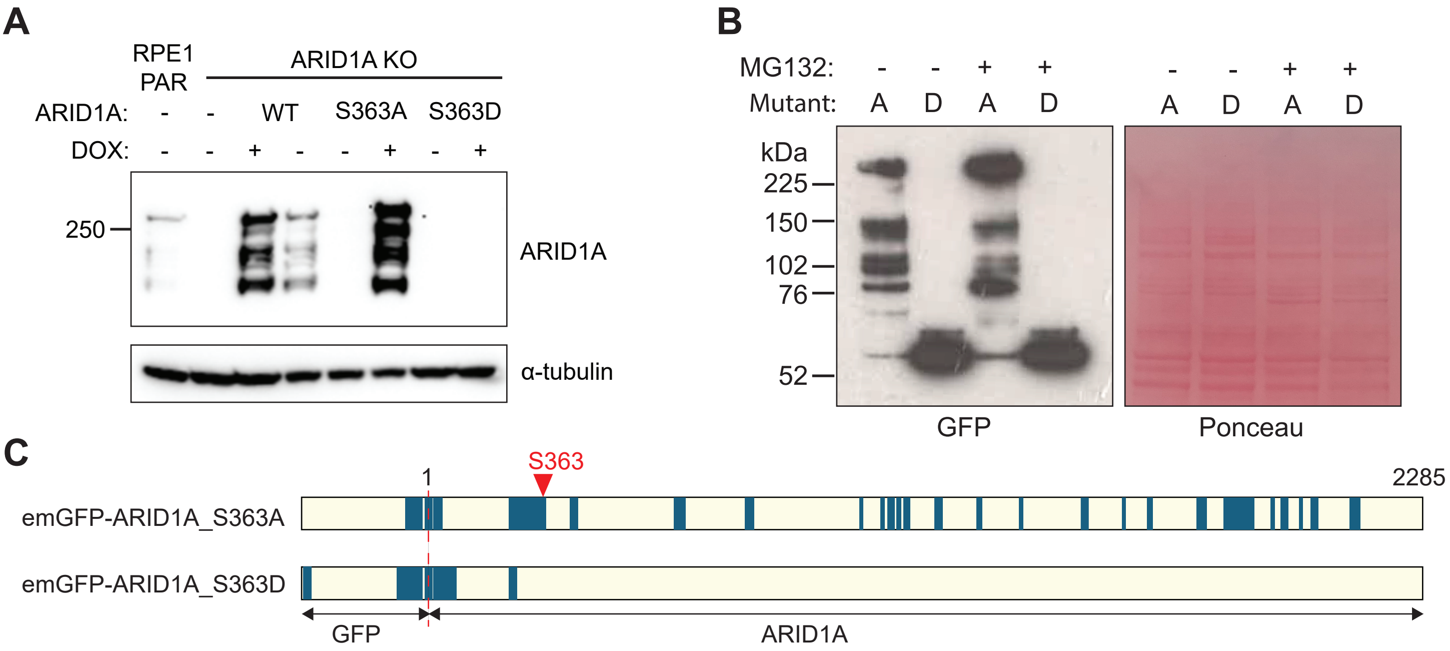


**D**

| Gene name | S363D | S363A |
|---|---|---|
| emGFP-ARID1A | 23 | 45 |
| SMARCC1 | | 8 |
| SMARCC2 | | 5 |
| SMARCA4 | | 3 |
| SMARCB1 | | 3 |
| ACTL6A | | 3 |
| SMARCE1 | | 1 |
| BCL7C | | 1 |
| SMARCD1 | | 1 |
| DPF2 | | 1 |
| SMARCA2 | | 1 |

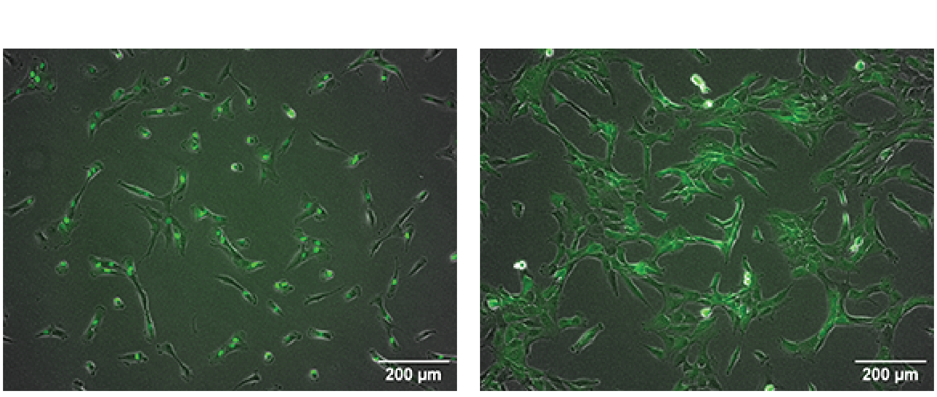


**Figure S4. Characterisation of ARID1A S363A and S363D mutants**. **A**, Immunoblot analysis of ARID1A in wt emGFP-ARID1A, emGFP-ARID1A_S363A, emGFP-ARID1A_S363D and parental RPE1 cell lines after 24 h induction with 1 ug/ml doxycycline. α-tubulin was used as loading control. **B**, Immunoblot analysis of emGFP-ARID1A fusion proteins in mutant ARID1A cell lines, untreated or treated with proteasome inhibitor MG132 (10 μM) for 12 h after 12 h doxycycline induction. Ponceau Red staining was used to assess equal loading. **C**, Primary sequence analysis of GFP-ARID1A mutant fusion proteins characterised by GFP affinity purification followed by MS analysis. Expression of ARID1A_S363A and ARID1A_S363D was induced for 6 h. Dark segments represent peptides detected by MS. **D**, BAF complex subunits identified in GFP immunoprecipitates from ARID1A S363A and S363D mutant cell lines. Numbers of peptide spectrum matches for each protein are shown. **E**, Fluorescence images showing the localisation of ARID1A S363A and S363D mutants in RPE1 cells after 24 h induction. Images of GFP-expressing cells were taken using an Evos M5000 (Thermo) at 10x magnification
